# Representational drift as the consequence of ongoing memory storage

**DOI:** 10.1101/2024.06.25.600729

**Authors:** Federico Devalle, Licheng Zou, Gloria Cecchini, Alex Roxin

## Abstract

Memory systems with biologically constrained synapses have been the topic of intense theoretical study for over thirty years. Perhaps the most fundamental and far-reaching finding from this work is that the storage of new memories implies the partial erasure of already-stored ones. This overwriting leads to a decorrelation of sensory-driven activity patterns over time, even if the input patterns remain similar. Representational drift (RD) should therefore be an expected and inevitable consequence of ongoing memory storage. We tested this hypothesis by fitting a network model to data from long-term chronic calcium imaging experiments in mouse hippocampus. Synaptic turnover in the model inputs, consistent with the ongoing encoding of new activity patterns, accounted for the observed statistics of RD. This mechanism also provides a parsimonious explanation for the diverse effects of experience on drift found in experiment. Our results suggest that RD should be observed wherever neuronal circuits are involved in a process of ongoing learning or memory storage.

---

The synaptic hypothesis of learning and memory postulates that memories are stored in the structure of the synaptic weight matrix in neuronal circuits ^1^. Theoretical studies have shown that network models built on this synaptic principle can store large numbers of patterns as fixed point attractors ^2–4^. However, with biologically constrained synapses, new learning implies the overwriting of previously stored memories ^5,6^. In the context of stimulus-driven neuronal activity, this overwriting would result in a distinct network response at two different points in time, even if the input pattern remained unchanged.

The advent of technologies allowing for stable, long-term recordings in awake behaving mice has indeed revealed that the neuronal activation underlying certain behaviors can change dramatically, even when sensory inputs do not. This phenomenon was first described in detail in the context of spatial memory in rodents ^7^. It was already known that in area CA1 of the hippocampus a unique pattern of place-cell activity quickly emerged upon exploration of a novel space ^8–12^, and was reliably re-evoked when the animal was returned to the familiar environment ^13–15^. However, the seeming stability of the hippocampal code only held true on relatively short time scales. Indeed, in familiar environments, both place-cell- and non place-cell activity slowly changed over days and weeks ^7,16–23^. These long time-scale changes in the neuronal code, dubbed representational drift (RD), have also been seen in other cortical areas, such as parietal, piriform, visual and auditory cortex ^24–28^. The phenomenology of RD is fundamentally similar in all of these cases: the identity of active neurons changes from session to session, although the sparseness of representation is stable. Furthermore, there is considerable heterogeneity in the stability of cells, or how often they take part in the code. It is not known what the mechanism generating the observed drift is, nor what its potential functional role might be.

Several computational studies have shown that ongoing plasticity can result in changes in neuronal dynamics reminiscent of RD at the population level ^29–32^. In these studies the plasticity acted as a source of noise, driving changes in the representation of already stored patterns. In fact, it was previously hypothesized that such changes might provide the substrate for a time-stamp of a given memory ^16^. Alternatively, it has also been hypothesized that RD may occur due to plasticity related to the encoding of new memories ^25^. From this perspective RD would be the unavoidable signature of ongoing memory storage due to overwriting ^5,33,34^. However, it remains unclear to what extent the mechanism of RD as ongoing memory storage is consistent with experimentally observed data. We therefore developed a biologically plausible network model which could be fit to data quantitatively, and which specifically allowed us to address the role of ongoing learning.

Through fitting of our network model to experimental data ^16^, we inferred that changes in neuronal activity from session-to-session are inherited from changes in the afferent inputs to cells in CA1. These changes are consistent with the large degree of synaptic turnover observed through Ca2+ imaging of dendritic spines in CA1^35^. Interestingly, the inferred synaptic turnover was random and uncorrelated from cell-to-cell, and followed simple Gaussian statistics. Although the types of memories which the hippocampus stores are, in general, highly correlated, an efficient means of data compression is to store just the differences between like memories, which are uncorrelated ^36–40^. Indeed, we show in our network model that the ongoing storage of such decorrelated patterns can account for the synaptic turnover, which, in turn, is responsible for RD.

This same mechanism provides a parsimonious explanation for a recent finding on RD in CA1 place cells. While RD in the spatial tuning of these cells depended on the amount of time spent exploring the environment under study, RD in overall rates depended only on the absolute passage of time^41^. In the context of memory storage, when the animal is engaged in the task, plasticity can readily occur between the active CA1 place cells and their presynaptic inputs, affecting CA1 place cell tuning. On the other hand, episodes unrelated to experimental sessions will not engage the same subset of place cells and changes due to plasticity will therefore be largely spatially untuned (in the experimental environment). The number of encoded episodes is simply proportional to the total time elapsed. In this way, RD in neuronal firing rates occurs as a function of time, while RD in neuronal tuning is biased more strongly towards time spent exploring, when the relevant assemblies of tuned cells are active. Overall, we find that the statistics and phenomenology of RD are well predicted by theories of ongoing memory storage.

## Results

Ongoing memory storage in a network model with biologically constrained synapses leads to the partial overwriting of previously encoded patterns, Fig.1. When such a network is driven by external input, the resulting patterns of activity will therefore change over time, even if the input is stable, Fig.1b-c. Such a drop in correlation of network activity over time, or representational drift (RD), has been observed in the hippocampus and other cortical areas. However, despite the broad qualitative similarity between plasticity-related overwriting and RD, it remains unclear if synaptic plasticity alone can provide a comprehensive account of the detailed statistical characteristics associated with RD. To answer this question, we fit a network model to experimental data from chronic Ca2+imaging in mouse hippocampus ^16^ and sought to reproduce the observed RD via changes in synaptic connectivity consistent with ongoing memory storage. We found differential effects of the spatially tuned versus untuned components of RD in the data, see Fig.S1, in line with recent findings ^41^. Specifically, the large drop in the correlation of the population activity from one session to the next was due largely to rate effects alone, while drift in the spatial tuning occurred on a longer time scale.

**Fig. 1.**
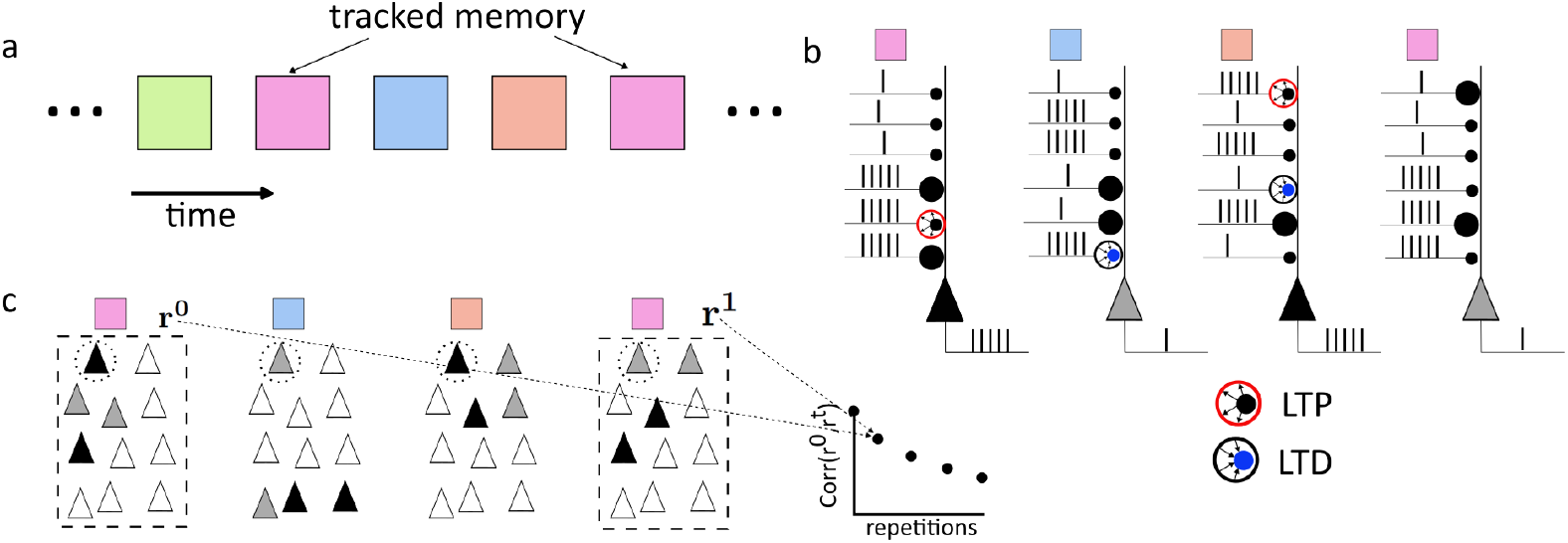
Ongoing memory storage generates representational drift. **a**. Memories are encoded in an ongoing fashion in time. Memory identity is indicated by color. We track and observe the neuronal activity corresponding to one memory in particular (pink square). **b**. Synaptic weights undergo plasticity as memories are stored. When pre- and post-synaptic activity is high, synapses may undergo potentiation (large red circle), while depression takes place if one cell fires strongly and the other only weakly (small blue circle). Active presynaptic cells are indicated by a train of action potentials. The postsynaptic cell fires strongly (black) or weakly (grey). Synaptic weights are strong (large circle) or weak (small circle). Note that due to the plasticity from the two intervening memories, the response of the post-synaptic cell to the same pre-synaptic pattern of activity, corresponding to the pink memory, has changed: its firing rate has decreased. **c**. The change in post-synaptic activity due to ongoing memory storage manifests itself at the population level as representational drift. Therefore, there is a drop in correlation between the initial pattern of neuronal activity given inputs corresponding to the pink memory *r*^0^ and the first repetition *r*^1^. If we assume that ongoing memory storage occurs between every repetition the correlation will continue to decrease.

### A spiking network model with synaptic turnover reproduces drift statistics in CA1

We sought to reproduce the observed RD in a network model in which plasticity occurred at different rates at spatially tuned versus non-spatially tuned synapses. Specifically, we modelled a local circuit of CA1 pyramidal and interneurons as sparsely-connected leaky integrate-and-fire neurons. Neurons in CA1 received input from area CA3 of the hippocampus and the entorhinal cortex (EC), modeled as Poisson neurons, Fig.2a (top). For simplicity we chose a fraction of CA3 cells to be place cells, while the EC inputs were spatially untuned. Allowing for weak to moderate tuning in the EC inputs did not alter the results qualitatively, see Fig.S4. We simulated the movement of a virtual animal along a circular track during a given session, Fig.2a (bottom). Large excitatory and inhibitory input currents dynamically balanced in the model ^42^, leading to the emergence of heterogeneity in firing rates and spatial tuning consistent with experimental findings ^43^, Fig.2b,c and Fig.S3a-g.

We modelled changes in the inputs to CA1 cells from session to session as a process of random synaptic turnover ^35^. Later we will show that this turnover is also consistent with the ongoing storage of memories. Specifically, we rewired a random fraction of the inputs for each cell in CA1, independently for the EC and CA3 pathways, Fig.2d. The rewiring fraction for CA3 was chosen to be smaller than that for EC, resulting in more gradual changes in place-cell tuning, in accordance with the data, Fig.S1. This synaptic turnover could lead to significant changes in the mean drive to CA1 cells from session to session, which is the mechanism responsible for RD, Fig.2e,f.

We chose the network parameters in order to closely match the statistics of RD observed in experiment. To do this, we first considered a simple statistical model for CA1 pyramidal cells which could be quantitatively fit to the data using standard least-square optimization, see Methods and Fig.S2. Inputs in this model were Gaussian random variables with zero mean, and with variance and temporal correlation optimized to fit RD statistics from the data. Input statistics in the full spiking network model were also Gaussian in the balanced regime, and could be calculated analytically, allowing us to map network parameters onto the statistical model, and hence fit the network to the data, see Methods and Fig.S3h. As a result, we were able to reproduce the drop in the Population Vector (PV) correlation seen in the data quantitatively Fig.2g, which takes both rate and tuning effects into account, as well as several measures of RD related only to active versus inactive cells, see Fig.S3i. The network model also qualitatively reproduced the gradual diffusion of place field location, Fig.2h, although the fraction of place cells was always larger in the network than in the data.

Finally, the mechanism of RD through synaptic turnover predicts that the likelihood of a cell to remain active on a subsequent session should be positively correlated with its firing rate. This is because high (low) firing rates are due to a large (small) number of excitatory inputs. Therefore, the rewiring of the same number of inputs is more likely to cause a low-rate cell to become inactive, than a high-rate one. We reanalyzed the data and found such a positive correlation, see Fig.S5. This is furthermore consistent with the greater stability observed in high-rate CA1 pyramidal cells across sleep sessions compared to low-rate cells ^44^.

### The synaptic turnover responsible for RD is consistent with ongoing memory storage

In the previous section we showed that synaptic turnover, modelled as a process of random rewiring of the synaptic inputs to CA1 pyramidal cells, could account for the statistics of RD observed in mice. At such a level of description, the mechanism behind the turnover itself remains unclear. We hypothesize that this rewiring process may, in fact, reflect the encoding of episodes or memories. In this scenario, much of the synaptic turnover, and hence RD observed from session to session, would be due to the storage of memories unrelated to the environment in which the recordings are made, Fig.3a-b. Specifically, instead of rewiring synapses randomly, we modified them according to a simple Hebbian plasticity rule. We imposed random binary patterns of activity for the CA3 and EC inputs, and the CA1 outputs, with sparseness *f* . We then potentiated synapses between co-active cells with probability *p*_+_ and depressed synapses between cells with differing activities with probability *p*_−_. These probabilities were different for the EC and CA3 inputs. We repeated this process many times, until the synaptic weight matrices from EC and CA3 to CA1 reached a statistical steady state, see Methods and Supplementary Information for more details. We then tracked one of the random patterns in particular, as if it were the experimentally observed activity pattern. In between observations of this pattern, we encoded other random patterns, the number of which we call the inter-session interval (ISI), Fig.3b. With this simple model we could calculate the temporal correlation of the inputs to CA1 cells analytically as a function of the plasticity parameters, and match them to the synaptic turnover process from the network, Fig.3c.

In our model, both synaptic turnover and RD are reflections of the changes in network structure due to the storage of memories, most of which are episodes unrelated to the ones observed during an experimental session. Nonetheless, given a Hebbian rule, plasticity occurring during a session, namely a repetition of the tracked pattern, can have an outsized effect on RD. This is because all of the neurons being tracked are co-active by definition, whereas other sparse, random patterns will have a small overlap with the tracked pattern. The repetition rate of a pattern therefore affects the degree of RD observed, although whether drift is increased or decreased depends on the details of the input statistics. In the following sections we show how this phenomenon can account for recent findings from both hippocampus ^41^ and piriform cortex ^25^.

### Ongoing memory storage accounts for the differential roles of time and experience on RD in place cells in CA1

Recent experimental work has shown that the spatially untuned component of RD in CA1 pyramidal cells occurs at a rate proportional to the absolute time elapsed between sessions on a linear track, while RD in the tuned component depends on the time spent exploring the track itself ^41^. The mechanism we propose here for RD suggests a potential explanation for this dissociation. Namely, we would expect that the ongoing storage of memories unrelated to the linear track itself would simply occur at a rate proportional to time. Such memories would naturally overwrite some synapses involved in the representation of the track, leading to RD. In principle, these synapses could be from spatially tuned or untuned presynatic inputs. However, it is reasonable to assume that it is the spatially untuned, contextual inputs which are more strongly shared between temporally proximate memories ^16^. On the other hand, it is only when the animal is actively exploring the track itself that the entirety of the spatially tuned inputs are active. Given an activity-dependent plasticity rule, we should thus expect maximal RD due to changes in the spatially tuned inputs to occur precisely during exploration.

In the framework of our network model, in which a series of patterns with spatial and non-spatial component are stored, the dissociation between time and experience is, in general, approximate. It is exact only in the limit in which the overlap between spatially tuned presynaptic inputs vanishes between patterns, Fig.4a. In this case, if we consider two sets of simulations in which one particular pattern (e.g. corresponding to the linear track) is repeated more often in one set than in the other, Fig.4b, then the RD in firing rates will depend only on the number of patterns presented, i.e. time, while the RD in tuning will depend only on the number of repetitions of the pattern of interest, i.e. experience. We ran simulations in which the encoded patterns consisted of a random fraction of spatial and non-spatial inputs, and hence the overlap in spatially tuned presynaptic inputs between pattern was not zero, but rather equal to the fraction of active spatially tuned cells *f*_*s*_ = 0.1. In this case, the RD in firing rates depended much more clearly on time that on repetitions, upper row of Fig.4c. The small discrepancy in the overlap was due to the fact that, all things being the same, the encoding of a random pattern in B led to a larger drop in correlation than a repetition of the tracked pattern in A. On the other hand, the RD in tuning depended both on time and experience, bottom row of Fig.4c. However, this result was true only when the patterns of presynaptic activity were stable over time. In fact, place cell activity in CA3 itself undergoes RD ^45^, suggesting that those presynaptic inputs should themselves vary from one repetition to the next, Fig.4d. Including this effect tipped the balance of spatial RD from time to experience through a combination of presynaptic RD and plasticity, Fig.4e. Interestingly, it also eliminated the small discrepancy in the rate correlation as the encoding of a random pattern or a repetition now led to similar drops in correlation.

### Ongoing memory storage accounts for the effect of repetition rate on RD

Recent experiments have revealed that the degree of RD in the piriform cortex of mice is reduced by increased repetition of familiar odors ^25^. This was shown by familiarizing two groups of animals with a particular odor over many sessions, and then repeating that same odor each subsequent session for cohort A, while cohort B was exposed to the familiar odor only after a number of sessions without odor, Fig.5a.

As was the case in the previous section, we again assumed that RD for the familiar odor was largely due to the storage of other patterns in piriform cortex, which occur in between sessions, see “random pattern” in Fig.5a. This assumption was enough to provide a simple explanation for the experimental finding, Fig.5b. Namely, if we consider the synaptic weight matrix for cohorts A and B, they were both initially identical due to the familiarization process. The storage of other patterns, which were random and uncorrelated with the familiar one, degraded the structure in the matrix which was correlated with the familiar pattern. However, in cohort A, the familiar pattern was repeated at the subsequent session, once again boosting the structure in the matrix, while in cohort B this was not the case.

We simulated this process using the same Hebbian plasticity rule as in previous sections, but with only a single input layer. We presented a pair of sparse, binary input and output vectors repeatedly until the synaptic weight matrix reached a steady state. We then repeatedly encoded the pattern to be tracked and considered this to be the familiarized state, see Methods for details. To simulate cohort A, we then encoded a number of random patterns in the network equal to the inter-session interval (ISI) after which time we again presented the familiar pattern, Fig.5c (black line). For cohort B we presented repetitions of the familiar patterns only after every eight repetitions in cohort A, leading to significantly reduced output correlations, Fig.5c (green line). The size of the difference in output correlations (RD) was strongly affected by the ISI, Figs.5d,e.

In the same set of experiments, an unfamiliar odor was presented to both cohorts with the same repetition rate and no difference in drift rate was found. We can account for this by running the same simulation as before, but now tracking both the familiar pattern, and one pattern uncorrelated with the familiar one, Fig.5f. This amounts to considering the structure of the weight matrix, and the output vector, with two distinct orderings. As long as the familiar and unfamiliar patterns are uncorrelated, the drift rate of the unfamiliar pattern depends only on the repetition rate and ISI, which were the same for both cohorts, Fig.5g-h. Finally, the familiar pattern with high repetition rate had a significantly reduced drift rate compared to the other three cases, in agreement with experiment, Fig.5i.

### Ongoing memory storage can explain diverse effects of experience

It may at first seem paradoxical that increased exposure to a spatial environment would lead to more RD in CA1 of hippocampus ^41^, while increased exposure to a familiar odor would actually reduce RD in piriform cortex ^25^. In the context of ongoing memory storage through Hebbian plasticity, both scenarios are possible, Figs.4 and 5, suggesting that this single mechanism provides a potential explanation.

We first note that the repeated storage of the *same* pattern tends to increase the correlation of the synaptic weight matrix with that pattern, Fig.6a-c. A period of familiarization provides a substrate for enhancing the effect of this repetition rate. The degree of this enhancement of correlation, and hence reduction in RD, depends strongly on the fraction of synapses which are updated to store each memory (learning rate). We quantified this by varying the number of times *k* a pattern was repeated before time *t* = 0. For *k* = 1 the pattern was novel. When the learning rate is small, increasing *k* leads to a build-up of correlation which was large compared to a single presentation, Fig.6a, whereas for large learning rates this difference was smaller, Fig.6b. Large *k* implies a weight matrix which is entirely correlated with the tracked pattern, and hence has lost all correlation with previously stored patterns, an unreasonable limit for an actual physiological memory system. However, the difference in RD for different repetition rates is already present for small values of *k* and even reaches a maximum for intermediate values when learning rates are small, black lines Fig.6c. These are the parameter values used to reproduce the findings from piriform cortex in Fig.5. The effect of varying the coding sparseness *f* is shown in Fig.S6.

When the input pattern is *not* the same from one repetition to the next then the effect of repetition rate is not as straightforward. We modeled this by keeping a fraction *s* of the active units in the tracked pattern the same from repetition to repetition, while choosing the other 1 − *s* randomly, Fig.6d. When *s* = 1 repetitions reduced RD as explained above. For *s* = 0 the pattern was effectively random and uncorrelated from one repetition to the next. For intermediate *s* the correlation of the *r*^*th*^ repetition with the original input pattern decreased as *s*^*r*^ which could lead to RD *increasing* for sufficiently high repetition rates, see Fig.6e. All other network parameters being the same, if RD was measured after a fixed number of repetitions, then there was a critical value of *s* below which repetition rate increased RD, Fig.6f.

## Discussion

Here we have argued that RD is, in large part, due to the storage of memories unrelated to the experiment in which RD is observed. Such a storage process implies changes in the synaptic weights of the network, which will necessarily alter its response to stimuli on subsequent sessions, even if upstream neuronal activity is unchanged. We have shown that the statistics of RD in CA1 of mice are well captured by a process of synaptic turnover in which inputs undergo rewiring from one session to the next. The fact that this synaptic turnover is consistent with the ongoing encoding of activity patterns to us strongly points to this as the underlying mechanism. After all, episodic memory formation, a major function of the hippocampus, involves precisely this kind of ongoing storage of information related to experiences during daily life. Ongoing memory storage can furthermore account for the differential effects of time versus experience in RD in CA1, as well as the effect of repetition rate, both stabilizing as found in piriform cortex and destabilizing, as in CA1, depending on the input statistics. These nontrivial and seemingly paradoxical findings cannot be explained by a process of random synaptic turnover, but are reproducible in a network model in which patterns are encoded via a Hebbian plasticity rule. Specifically, whether experience led to more or less drift in our computational model depended on the stability of the input patterns. This result is consistent with the finding that CA3 input patterns themselves drift ^45^, while representations in the olfactory bulb, the primary input to piriform cortex, are extremely stable over time ^46^. More generally, the fundamental mechanism here is the *interference* between stored patterns. Precisely how this interference manifests itself at the level of RD will depend on the correlation between these patterns. Here, for simplicity, we considered all patterns to be random and uncorrelated.

Other computational modeling work has also proposed that RD may arise due to plasticity-driven perturbations ^29–31^. In ^29^ the authors studied a network with prescribed synaptic weights which allow for the storage of a large number of sequential patterns. They showed that randomly perturbing the synaptic weight matrix generates RD while maintaining robust sequences. RD can also arise in a spiking network with a symmetric spike-timing dependent plasticity rule, coupled with a homeostatic mechanism ^30^. Specifically, if the initial network structure (synaptic weight matrix) exhibits clustering, ongoing plasticity allows for individual cells to leave their cluster and join a new one, all the while maintaining the clustered structure at the network level. RD occurs in a similar fashion in networks which minimize the mismatch between the similarity of pairs of input patterns and the corresponding pairs of output patterns (Hebbian/antiHebbian networks) ^31^. Namely, ongoing plasticity allows the network to explore the degeneracy in the solution space by undertaking a random walk along the manifold of equally optimal output patterns. An important conceptual difference between our work and previous studies is the nature of the synaptic turnover itself. While those studies showed that *already-stored* patterns of activity undergo RD in the face of ongoing plasticity, we ascribe the plasticity to the encoding of *new patterns*. Namely, we have shown that RD is consistent with the inevitable interference between patterns when learning occurs, and hence not necessarily just a consequence of noise once learning is done. The mice in the experiments we have studied are not exposed to explicit, task-dependent learning between sessions. Rather, the “learning” process may simply be the storage of episodes, unrelated to the exploration. The hippocampal circuit plays a central role in this type of memory ^47,48^, while there is evidence that piriform cortex participates in olfactory associative memory formation ^49^.

*The stability paradox* We have sought here to provide a plausible network mechanism for RD, and have not addressed the fundamental paradox of how to maintain stable behavior in the face of such neuronal instability. In fact, drift does not appear to adversely effect the performance of mice in a variety of memory-dependent tasks ^24,25,50^. Several previous studies have addressed how this might be possible. Firstly, it has been hypothesized that there may be a low-dimensional manifold which represents the task-relevant projection of the population response, and which is invariant to RD ^27,51,52^. In this scenario many distinct patterns of the high-dimensional population activity can have the same projection on the relevant, low-dimensional manifold. If RD does not affect this projection, i.e. it is constrained to the “null-space” of the manifold, then task-relevant variables can be stably read out. While neuronal representations undergoing RD do appear to be low-dimensional, the direction of drift has not been found to be orthogonal to this manifold in general ^7,25,53^. Alternatively, despite ongoing changes, the patterns of neuronal activity observed in-vivo retain some significant correlation from session to session ^7,16,21,24,25^. Therefore, decoders trained on a given session will perform above chance for subsequent sessions, albeit with degraded accuracy. In any case it remains unclear precisely how behavior remains unaffected by the observed drift. One potential solution to this is to allow for compensatory plasticity in downstream circuits so as to stabilize the readout performance ^30,51,53–55^. Another is to seek some higher-order population structure which remains stable in the face of ongoing RD ^19,21,22,27,51,52^.

Yet despite recent theoretical advances and experimental findings, the stability paradox remains. It may be that behavior depends on a neuronal representation which is distributed across several cortical areas, and that RD is greatly reduced in some of the them compared to those observed up until now. In the case of the hippocampus, it is hypothesized that memories are transferred to higher-order cortical areas for long-term storage via a consolidation process which can take weeks, months or years depending on the species ^56–59^. Computational models of distributed memory systems in which fast learning (and hence fast forgetting) circuits drive plasticity in slower-learning downstream circuits in a multi-layered framework, show qualitatively enhanced memory capacity compared to single-area models ^34^. Such models leverage a hierarchy of time-scales of synaptic plasticity to allow for both fast encoding as well as long lifetimes ^6,34,60^. A hallmark of a model in which the timescale of plasticity is spatially distributed across cortical areas, is a concomitant array of timescales for RD. Only future experiments in which the activity of neuronal populations across several cortical areas are recorded simultaneously over long periods of time will reveal if RD acts on distinct timescales in different brain areas.

## Methods

### Data analysis

We analyzed previously published data ^16^. In the experiment, mice repeatedly explored two familiar environments (one in the morning, the other in the afternoon) over the course of two weeks, with imaging sessions every other day (8 total sessions). We separately analyzed leftwards vs rightwards running epochs for each environment, and then pooled the data together. In total, we analyzed data from 5 mice. Of the five mice, two have only one spatial map for each environment, while three have multiple spatial maps for each environment, as previously analyzed ^20^. Unless otherwise specified, all the analysis was performed separately for each map, and then results were pooled together.

Calcium events and place field maps were extracted as in the original paper ^16^. Briefly, the linear tracks were divided into 24 bins, each 4cm in length. For each spatial bin, the total number of events in each session was extracted, together with the total time-occupancy of each bin. The event rate map is then calculated by dividing the total number of events per bin by the bin occupancy. The two bins at each extreme of the track were excluded from the analysis to limit reward delivery effects.

In Fig.S1, the rate maps follow two different types of normalization: in Fig.S1a, the event rate map for each cell was normalized by the maximal firing rate over all sessions. In Fig.S1c, the event rate in each session/bin as normalized according to the maximal value within each session.

### Rate and tuning correlation

To compute rate correlations across sessions, we defined a rate population vector **r**_**t**_ ∈ ℝ^*N*^ . *N* was the total number of neurons, and each entry of the vector was the mean firing rate of each cell in session *t*. We then computed the Pearson correlation coefficient to quantify the rate similarity between two sessions, as in Fig.S1d-f.

To quantify the similarity of spatial tuning across session, we defined a Tuning population vector of length *N* × *n*, where *n* is the number of bins of the linear track. The vector then contained the rate of each cell in each spatial bin, normalized in such a way that the sum of the rates for each cell over the different spatial bins was constant in all sessions. We then computed the Pearson correlation coefficient of such vectors for pairs of session to quantify the tuning similarity. The procedure ensured that changes in mean firing rates of cells from one session to the next would not affect the tuning correlation.

### Place fields

To be considered a place cell in a given session, we employed the following procedure. First, for each cell, we computed the spatial information per spike as :

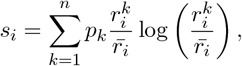

where *i* was the cell index, and the sum runs over all *n* spatial bins, 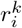 was the event rate of cell *i* in bin *k*, 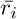 was the mean event rate over all bins, and *p*_*k*_ was the occupancy of bin *k* (the fraction of time spent in bin *k*). *s*_*i*_ was then the spatial information (measured in bits/event) of cell *i*. Then, we generated surrogate data by shuffling the position of the animal with respect to the time of the calcium events, and calculated the spatial information for each cell and each shuffle. We then compared the value of the spatial information of each cell to the null distribution generated with the surrogate data. If the value was larger than the 95th percentile of the null distribution, than the cell was defined as a place cell in that session.

The place field width of each cell was defined as the number of contiguous bins where the event rate is larger than 50% of its maximal value over all the bins.

### The statistical model

In order to fit the network model to data in Fig.2, we first fit a simple statistical model to the data, and then mapped the network parameters onto the resulting fit parameters as a starting working point. For the statistical model we took cells in CA1 as binary units which received inputs from two sources, CA3 and layer III of entorhinal cortex EC. The total input to a cell *i* at time *t* (time measured in sessions) from CA3 was written 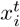 and from EC 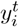. Both inputs were Gaussian random variables with zero mean and variances 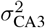and 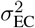 respectively. A neuron *i* was active at time *t* if its total input 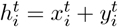 exceeded a threshold *θ*, and was otherwise silent. Specifically, the activity was written as 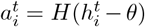, where *H*(*x*) = 1 if *x >* 0 and *H*(*x*) = 0 if *x* ≤ 0 is the Heaviside function.

To model RD we allowed for the inputs to change over time according to:

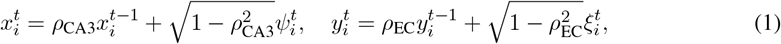

where the autocorrelations *ρ*_CA3_ and *ρ*_EC_ ∈ [0, 1] and 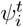 and 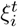 were Gaussian random variables with mean zero and standard deviation *σ*_CA3_ and *σ*_EC_ respectively. The update rules Eqs. (1) ensured that the input distributions were stationary. Inhibition is implicitly assumed to have the effect of subtracting the mean of both inputs so that they are centered at zero.

The statistical model had four parameters once we rescaled them by the standard deviation of the inputs from EC: the ratio *σ*_CA3_*/σ*_EC_, the rescaled threshold *θ/σ*_EC_ and the autocorrelations *ρ*_CA3_ and *ρ*_EC_. Note that the input dynamics Eqs. (1) could be formulated in continuous time as an Ohrnstein-Uhlenbeck processes, as shown in the Supplementary Material . The continuous formulation allowed us to calculate the time constant of the decay in correlation of the inputs analytically, yielding 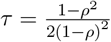 .

According to the definitions above, the state of the network was defined by a vector 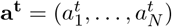 where *N* was the total number of neurons. *Fraction of active cells:* The fraction of active cells can be calculated analytically for the statistical model. The probability that a cell with a given input *y* is active can be written Pr(*y > θ* − *x*). Integrating this probability over all possible values of *y* gives the likelihood of any cell to be active, or the fraction of active cells, *f*_*a*_. This fraction is therefore

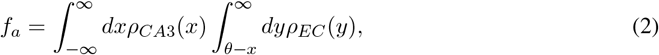

where 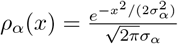

### Fit of the statistical model

To fit the statistical model, we only considered mice with a single map per environment (two mice in total), since with multiple maps the distribution of the number of sessions in which each cell was active and the survival fraction are not well defined (with multiple maps not all maps are visited in all sessions). We considered data from these two mice (N=1649 total recorded cells). The statistics used to fit the model were the distribution of sessions each neuron is active, the survival fraction (probability an initially active cell continued to be active on subsequent sessions), the population activity overlap between different sessions, and the fraction of active cells in each session, see Extenden Data Fig.2. To fit the data, we adjust the four free parameters of the model using least-square optimization. Specifically, to produce Fig.S2g-i, we discretize the (*ρ*_*CA*3_, *ρ*_*EC*_) space on a 20 × 20 grid, and for each position on the grid, run the Scipy ^61^ implementation of a Basin-hopping optimization algorithm minimizing the sum of the squared residuals between numerical simulations of the statistical model, and the experimental data. For the numerical simulations of the model, we consider *N* = 20000 neurons.

### Network model description

CA1 was modeled as a network of integrate-and-fire excitatory (E) and inhibitory (I) neurons with I-I, E-I and I-E connections, but no recurrent excitation, as prescribed by anatomical constraints. CA3 cells are modeled as Poisson neurons a fraction of which were spatially modulated. They projected onto both excitatory and inhibitory CA1 neurons. Additionally, CA1 neurons received excitatory inputs from a layer of non-spatial Poisson neurons (from layer III of EC). Detailed equations and parameters are given in Supplementary Information.

### CA3 place fields and non-spatial inputs

CA3 neurons were modeled as Poisson neurons. In any given environment, a fraction *f*_*CA*3_ of CA3 neurons were active. Of the active neurons, a fraction *f*_*s*_ of the population had a spatially modulated firing rate, while the remaining fraction had a constant firing rate. For simplicity, we considered a ring topology, so that the spatial position of a virtual animal was parametrized by an angle *ϕ* ∈ [−*π, π*]. The firing rate of spatially selective neurons was modulated according to a Von Mises distribution:

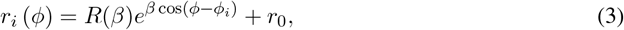

where *ϕ*_*i*_ was the center of the place field of neuron *i, r* _0_ a baseline firing rate, and 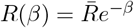 where 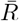 was a constant that set the maximal firing rate. The parameter *β* determined the sharpness of the place fields.

The layer of neurons providing non-spatial inputs were modeled as Poisson neurons with constant firing rate ν. Also for this layer, only a fraction *f*_*EC*_ of the population was active in any given environment.

### Connectivity matrices

Connectivity matrices for all subtypes of connections within the CA1 populations were random and sparse. Each neuron had a probability of connection to other neurons in the respective subpopulations equal to *α*_*i*,*l*_ = *K*_*i*,*l*_*/N*_*l*_, where *N*_*l*_ was the number of neurons of the *l*th population, *l* ∈ {*E, I*}, and *i* ∈ {*E, I*} . If not specified otherwise, we fixed the in-degree of CA1 pyramidal cells from interneurons to be fixed and equal to *K*_*I*_ (all pyramidal cell receive the same amount of inhibitory inputs). On average, each CA1 cell received projections from *K*_*i*,*{E*,*I}*_ neurons. The connectivity between the layer of non-spatial Poisson neurons and CA1 is also random and sparse with connection probability *α*_*EC*_. In the main text, we consider a uniform connection probability also for the CA3 → CA1 projections; in the Supplementary Information we discuss how to consider a phase bias in such connections.

Throughout, we assume all connection probabilities are the same and equal to *α* = 0.125.

### Input correlations in the network model

To implement the session-to-session changes in spatial and non-spatial inputs provided by Eqs. (1) in the network model, we needed to calculate the autocorrelation of inputs where changes may occur either due to changes in the input firing pattern, or in the connectivity matrices themselves. The general expression for such inputs is

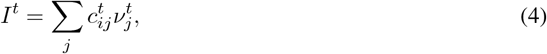

where 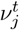 is the firing rate of the input neuron *j* at time *t*, 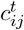 is the connectivity matrix element at time *t*, and we omitted conductances and time constants. In the Supplementary Information we provide detailed calculations. The expression one finds for the autocorrelation is

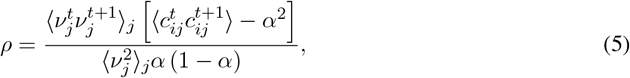

where *α* is the connection probability to the input layer. In general then, the level of correlation depends on the correlation in the input firing patterns, and on the degree of synaptic plasticity. Note that in order to obtain completely uncorrelated inputs from one session to the next, one must have 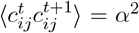, which implies 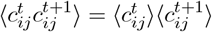, i.e. complete rewiring from one session to the other. Assuming completely uncorrelated input firing patterns from one session to the next results in:

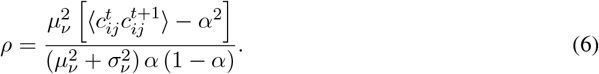

On the other hand, if we assume that the input firing patterns are the same over sessions, we have:

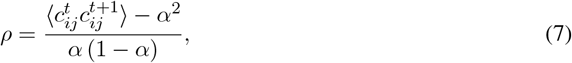

which, as shown in the Supplementary Information, can be written as:

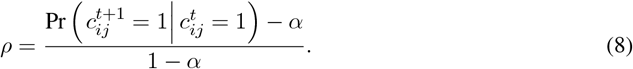

In the following, for both spatial and non-spatial inputs, we assumed that the input firing rates were constant from one session to the next. In this case, the fraction of connections rewired from one session to the next depending on the autocorrelation *ρ* is shown in Fig. S 3h.

### Variance of spatial and non spatial inputs

The average current from either the spatial or non-spatial inputs layer had the form:

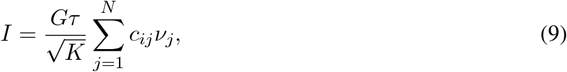

where we have scaled the synaptic weights as 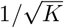. The expected value of the input is therefore

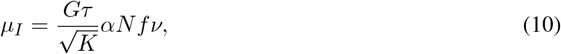

where *α* is the connection probability, *f* is the fraction of active pre-synaptic cells, and ν is their mean rate. If we define the normalization factor as the mean number of active inputs *K* = *αf N*, we have 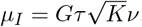. The variance of such current, using the results from the Supplemetary Information, was given by

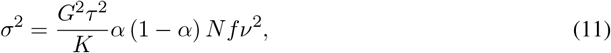

where we neglected the intrinsic variability of the Poisson process (which goes to zero as Δ*t*^−1^). Again using the definition of the normalization factor *K*, we have

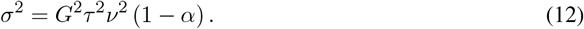

Given the variances ratio obtained fitting the statistical model 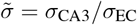, we can then fix the firing rates/synaptic weights of CA3 and EC inputs via the relation:

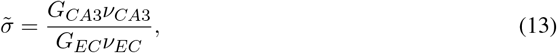

since both the time constant and the connection probabilities are the same for the two layers. We then fix ν_*CA*3_ = ν_*EC*_ (same average rate of the two layers), and set 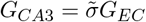.

### Fit of network simulations to data

In network simulations the virtual animal runs at a constant speed of *v* = 12 cm/s over a circular track of length *L* = 84 cm. The position of the animal on the track is parametrized with a phase *ϕ* ∈ [−*π, π*]. Analogously to the experiment, we simulate 8 sessions each consisting of 20 laps on the track (i.e. one session lasts 140 seconds of simulation time). For place fields analysis, the track was divided into 20 bins each of 4.2 cm length.

Given the analytical formulas derived in the previous section we were able to match both the variances of the input distributions as well as their autocorrelations to the values from the fit of the statistical model. The means of the input distributions were not zero, as in the statistical model, but the network operated in a balanced regime in which currents from inhibitory interneurons cancelled the mean excitatory drive to cells in the mean, leaving their membrane potential near threshold to spiking. Therefore, fitting the variances and correlations alone set the network at a working point in which the statistics of RD were close to the statistical model, and hence the data. We then made slight changes to parameters by hand in order to improve the fit. In order to fit the population vector correlation we needed to compare to the calcium event rate in the data. We did this by applying an exponential kernel with time constant *τ*_*c*_ = 500ms to the spike train from each cell in the spiking network. A calcium event was detected whenever the smoothed signal crossed a threshold *θ*_*c*_, and imposed a minimum inter-event interval of 500ms. The threshold value which minimized the mean squared error of the fit was *θ*_*c*_ = 0.16837.

**Figure 2:** The network model simulated in Fig.2 is that described above, with parameter values given in the Supplementary Information. RD is modeled by randomly rewiring a fraction of connections to each CA1 cell from the CA3 and EC inputs. Specifically, the probability of a synapse present at time *t* being removed is defined as 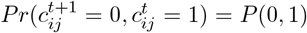, while the probability of a new synapse appearing is *P* (1, 0). We then define the rewiring fraction as the total fraction of altered synapses, i.e. 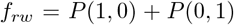, where there is a different fraction for each input source given by 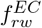 and 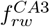. Using the definition 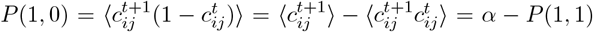, and with *P* (1, 0) + *P* (0, 1) and the definition of the temporal correlation, Eq.8, we find 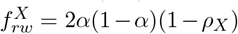. Parameters: *α* = 0.125 is the connection probability, and *ρ*_*CA*3_ = 0.95, *ρ*_*EC*_ = 0.35 are the temporal correlations of the inputs from the fit of the statistical model.

**Fig. 2.**
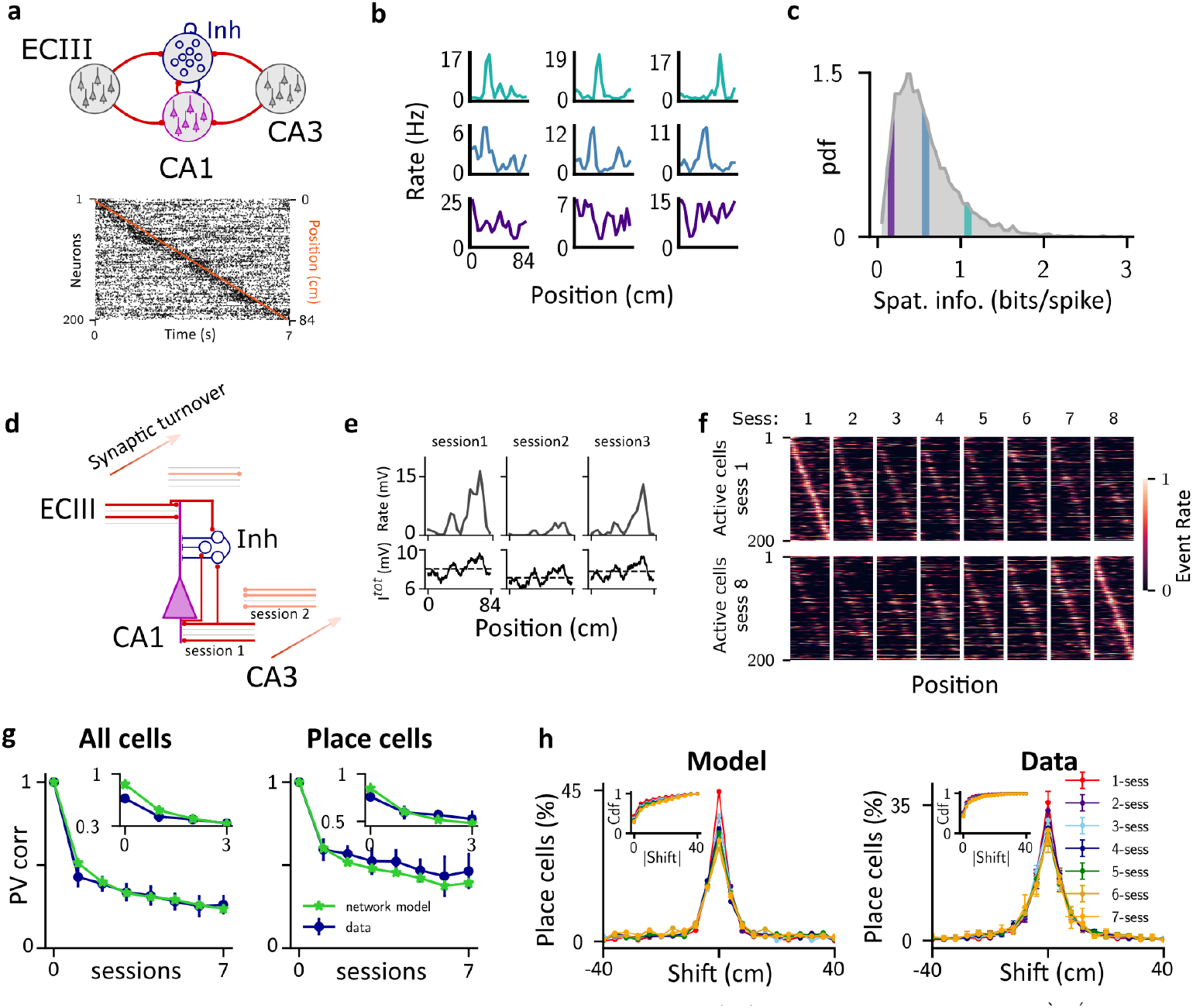
A spiking network model with random synaptic turnover reproduces drift dynamics. **a**. Network architecture and sample raster plot of CA1 pyramidal cells. **b**. Heterogeneous response profiles for sample CA1 cells. The color indicates the value of spatial information from c. **c**. Histogram of spatial information for all CA1 cells over one session. **d**. Synaptic turnover from session to session is modeled by randomly rewiring a fraction of inputs to each CA1 pyramidal cell, independently for EC versus CA3 pathways. **e**. Tuning curve (top row) and total input (bottom row) for one example cell over three sessions. The dashed line in the bottom panel indicates the average total input along the track for each session. **f**. Place field maps for 200 randomly selected active cells found on session 1 (top) or session 8 (bottom), ordered according to their place field positions. **g**. PV correlation of all cells (left), and only place cells (cells significantly spatially tuned in both sessions) (right). The insets show the first four points of the respective curves, where the initial point (within-session correlation) is computed considering odd vs even trials. **h**. Distribution of the centroid shift for different number of elapsed sessions (color-coded). Inset: cumulative distribution of the absolute shift. See Methods for model details and parameter values.

### Plasticity model

In the network simulations used to fit the experimental data in Fig.2 changes in inputs from one session to the next were modeled as a process of synaptic turnover, without specifying the precise mechanism responsible for this. One possibility is that this turnover is due to the storage of patterns in the network. Specifically, we assume that between sessions a fixed number of patterns are encoded, which we call the inter-session interval (ISI). Each pattern is a binary vector in which only a fraction of cells *f*_*s*_ are active. Of the active cells, we assume that a fraction *f* are strongly active, while the remaining 1 − *f* are weakly active. If we are modeling CA1, as in Figs.3 and 4, there would be three such vectors, while for Figs.5-6 there is only one input layer and hence only two vectors. To store a pattern we apply the following plasticity rule. If the pre- and post-synaptic cells are both strongly active, then the synapse is potentiated with probability *p*_+_. If one of the cells is strongly active while the other is weakly active then we depress the synapse with probability *p*_−_. If both are weakly active no change is made at the synapse. For the CA1 model, these probabilities can be different for CA3 versus EC inputs. Synapses are binary, as before, i.e. *c*_*ij*_ ∈ {0, 1} . To quantify RD we calculate the correlation in the output pattern at time *t* with that at time *t* = 0 given the same (tracked) input patterns. If the input patterns are the same, then RD can only be due to changes in the network connectivity. If the input patterns change, as is the case with CA3 inputs in Fig.4e, then RD is affected both by this as well as changes in network connectivity.

**Figure 3:** We mapped the Hebbian plasticity model onto the process of synaptic turnover in the network model by matching the decay in the correlation of the inputs a cell receives over time, see Fig.3c. The parameter values used to generate the lines in Fig.3c were: 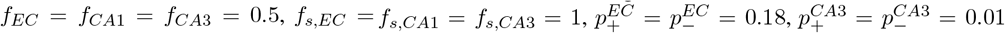, 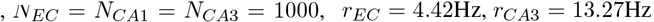 The circles were generated using the same rewiring process as that used to model synaptic turnover in Fig.2f. The only difference is that *N* = 1000 for all populations instead of 4000.

**Fig. 3.**
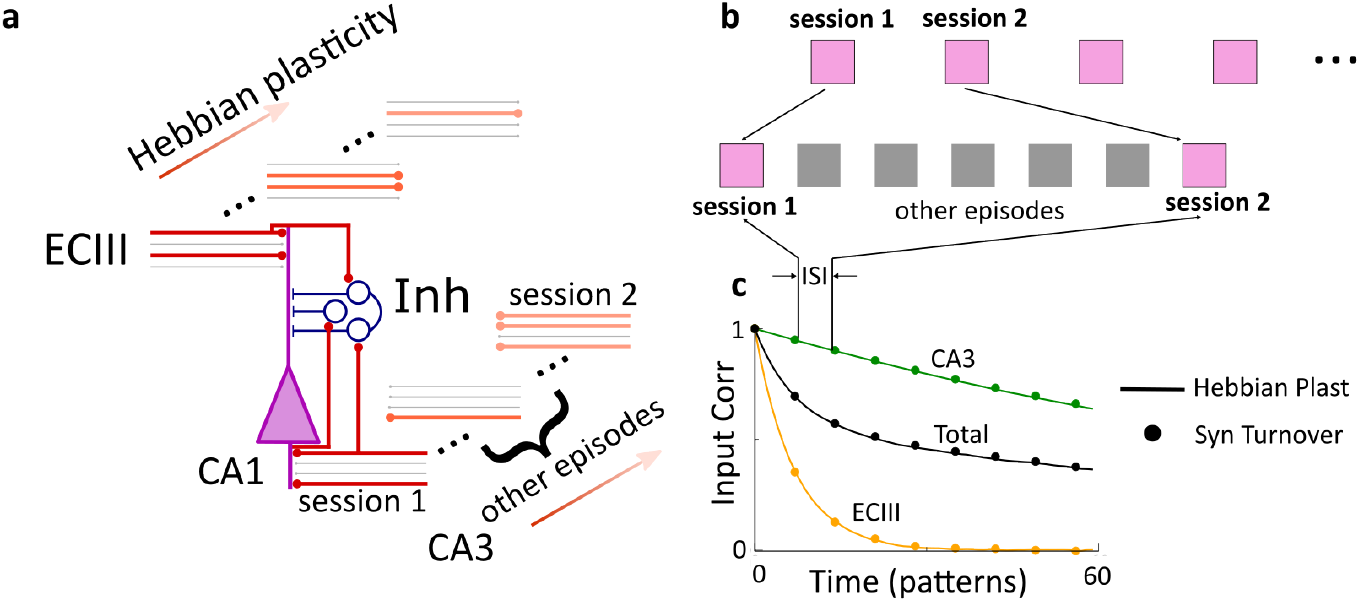
Synaptic turnover is consistent with the ongoing storage of random patterns. **a**. The synaptic turnover used to fit RD from the data is now generated through the encoding of random patterns with a Hebbian plasticity rule. **b**. We assume that in between sessions in which the activity patterns are tracked, there are a number of random patterns encoded. **c**. The learning process is fit to the synaptic turnover by matching the drop in correlation in input to CA1 cells over sessions. The circles indicate the values of correlation which correspond to the snapshots of activity shown in Fig.2f, see Methods for details and parameter values.

We also performed network simulations with the plasticity process and observed RD as in Fig.2, see Fig.S7. For those simulations we modeled a population of CA1 cells as linear threshold units, i.e. the activity of cell *i* was given by *r*_*i*_ = [*I*_*i*_ − *θ*]_+_, where *I*_*i*_ was the total input, *θ* was the threshold, and [*x*]_+_ = *x* if *x >* 0 and is zero otherwise. Active cells in EC had a constant firing rate *r*_*EC*_ while active cells in CA3 were spatially modulated according to a von Mises distribution as in Eq.3. The threshold was *θ* = 0.3 and for the von Mises rates the parameters were 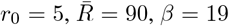. *r*_*EC*_ is chosen such that the ratio *σ*_*CA*3_*/σ*_*EC*_ = 1.16 in order to match the fit from the statistical and network models. The initial condition for the connectivity matrix was random with connections probability *p* = 0.33. We tracked the 100th pattern stored and plotted snapshots of the activity in CA1 starting at time *t* = 140 and every 7 time steps for eight “sessions”. Plasticity was not applied for additional repeats of the tracked pattern here. Neurons were considered inactive if their mean firing rate was less than 0.1.

**Figure 4:** We used the same network model of linear-threshold units as described above with the following parameter values: 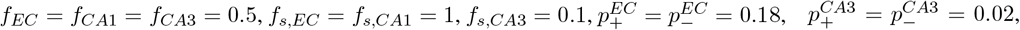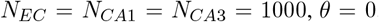. All other parameter values were the same. Drift in the CA3 inputs was modeled by allowing the position of the *i*^th^ place cell (center of the von-Mises distribution) at time *t* to shift with respect to the position at time *t* − 1 according to 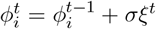, where *ξ* is uniformly distributed from {−*π, π*} and *σ* = 0.0447. Plasticity occurred according to the rule described above. To calculate the correlation of the tracked pattern in CA1 with the activity at time *t* = 0, the neurons in CA1 at time *t* were driven by the input pattern 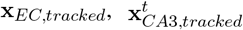.

**Fig. 4.**
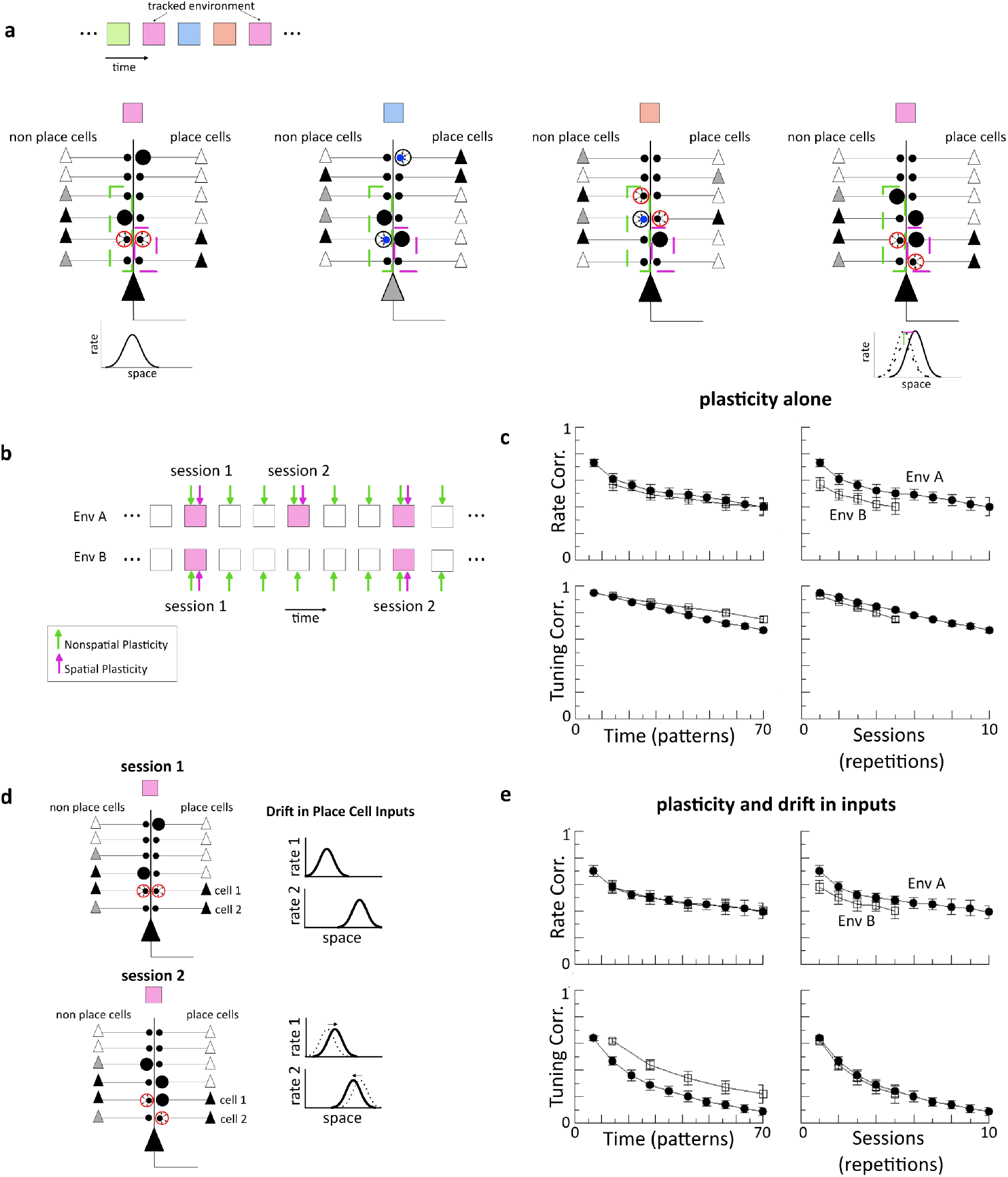
Ongoing memory storage is consistent with differential effects of drift on rate and tuning. **a**. We consider the storage of many memories, indicated here by color, all with spatial and non-spatial features, and one of which (pink) is tracked explicitly. Only changes to synapses which are active in that environment (dashed boxes in cell schematic) will generate observable RD. In the limit of sparse place cell coding the storage of the blue and orange memories readily causes drift in non-spatial inputs, but not in the spatial ones. As a result, RD in non-spatial features is proportional to the total number of memories stored, while RD in spatial tuning is due only to repetitions of the tracked memory. **b**. Simulation protocol for which the repetition rate for environment A is twice that of B. In the sparse spatial coding limit the rate correlation depends on total patterns stored (time) while the tuning correlation depends only on repetitions of the tracked pattern (session). **c**. Results of simulations when the sparseness of spatial coding *f*_*s*_ = 0.1. The drop in tuning correlation is affected both the number of repetitions as well as interference from other memories. **d.-e**. When spatially tuned inputs already exhibit drift, as has been observed in CA3 place cells, the combined effect of sparseness and drift results in the tuning correlation being dominated by the number of repetitions. See Methods for model details and parameter values.

**Figures 5:** To model the results from pririform cortex we studied a two-layer network. As before, patterns were random, binary and sparse with sparseness *f*_input_ = *f*_output_ = *f* . We stored many such patterns until the network reached a statistical steady state. We then tracked one particular pattern, for which the input vector can be written **x**. For simplicity we considered linear neurons, and hence the output pattern at time *t*, **y**^*t*^ = **C**^*t*^**x**, where **C**^*t*^ was the connectivity matrix. Simulations with linear-threshold neurons revealed that the nonlinear threshold did not qualitatively affect the results (not shown). To model the effect of familiarization we presented the same (tracked) pattern to the system *k* times before time *t* = 0 where *k* = 1 indicated a novel pattern. We quantified the RD by calculating the output correlation

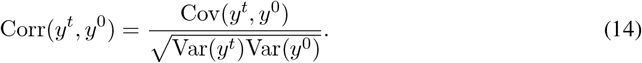

**Fig. 5.**
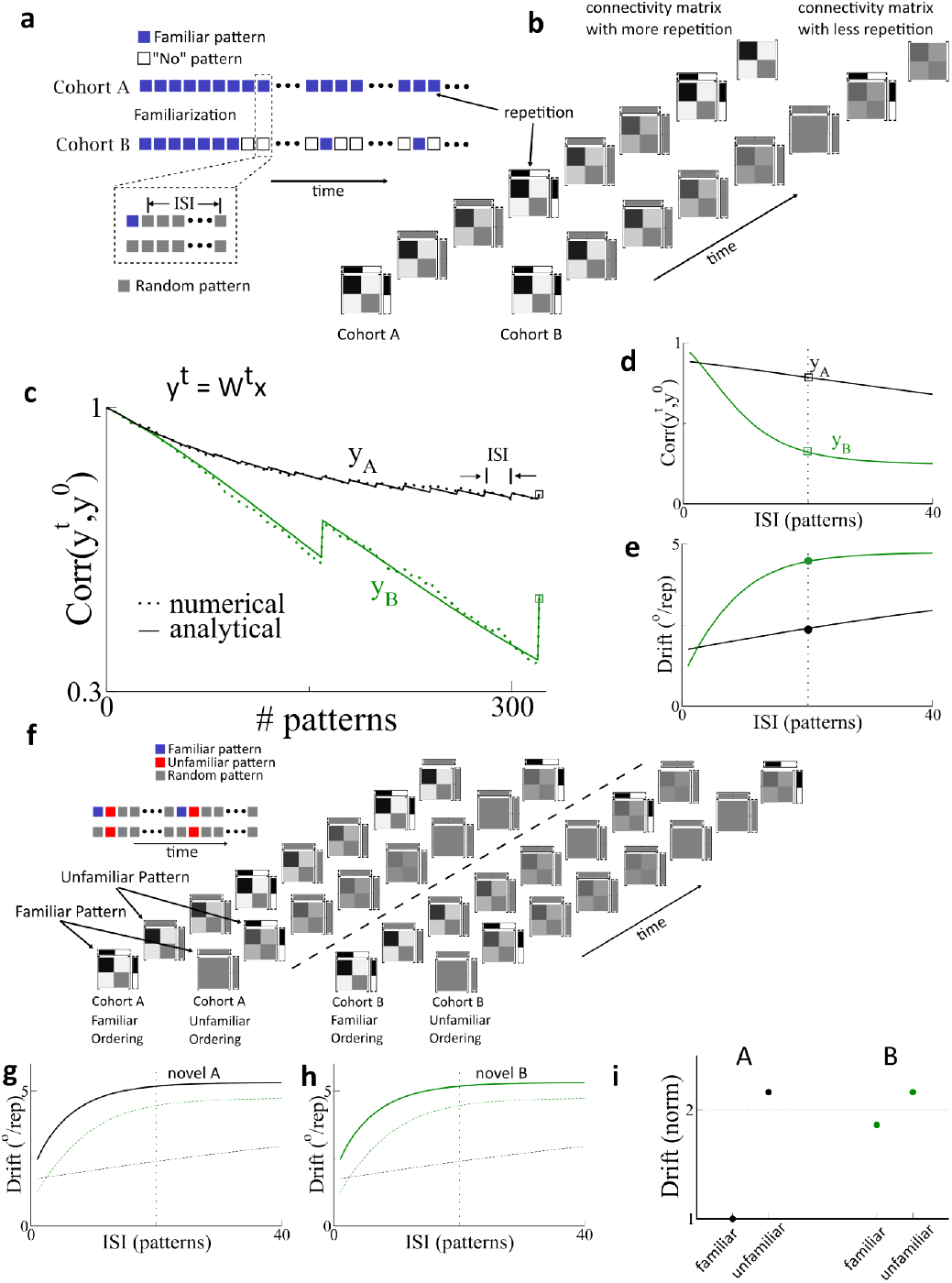
Repeated exposure to same input pattern reduces drift. **a**. Protocol for testing the effect of repetition rate on drift. Two cohorts are repeatedly presented a pattern (blue squares) during a familiarization period. After this time cohort A is presented the familiar pattern every session while B is presented the familiar pattern every eight sessions. Crucially, we assume that a number of additional patterns, uncorrelated with the familiar one are encoded between sessions (inter-session interval, ISI) (grey squares). **b**. Illustration of the effect of repetition rate on network connectivity. Repetition boosts network structure correlated with the familiar pattern, thereby reducing drift. **c**. Output correlation for familiarized pattern with a total of 16 repetitions for cohort A and *ISI* = 20, while the repetition rate for B was 8 times less. Dotted lines are from simulation of the network model with Hebbian plasticity while solid lines are the solution of the corresponding Markov process. **d.-e**. Output correlation and drift rate as a function of the ISI, i.e. the number of random patterns encoded between “sessions”. The vertical line indicates the value of ISI used in **c. f**. Illustration of encoding of familiar and unfamiliar patterns. Because the repetition rate is the same for the unfamiliar pattern for both cohorts, the resultant drift is also the same. **g.-h**. Drift rates for the unfamiliar patterns. **i**. Drift rates normalized by the familiar case for cohort A. Parameters: *p*_+_ = *p*_−_ = 0.02, *f* = 0.15, *N* = 1000. For the familiar cases the tracked pattern was encoded *k* = 5 times at time zero, whereas for the novel cases *k* = 1. See Methods for model details.

The variance and covariance terms can be expressed in terms of first and second-order statistics of the network connectivity, which in turn can be calculated as a Markov process. Specifically, the connectivity depends on transition matrices for the presentation of random patterns or repetitions of the tracked pattern. These matrices can be applied in any order to model a given protocol, see Supplementary Information for detailed calculations.

Conversely, instead of calculating the correlation over time we can calculate the drift, which we defined as

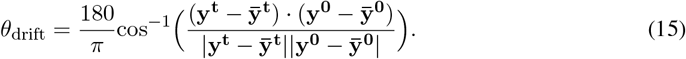

**Figure 6:** For Fig.6a-c this is exactly the same model as for Fig.5. In Fig.6d-f we consider input patterns which themselves also undergo drift. To model this, we assumed that the input *i* for the *r*^th^ repetition, 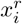 was identical to 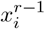 with probability *s* and was otherwise set active with probability *f*, and otherwise inactive with probability 1 − *f* . Therefore for *s* = 1 the input pattern was stable while for *s* = 0 the input pattern was random and uncorrelated from repetition to repetition. At the level of the Markov process, this meant that for each presentation of the tracked pattern we applied the transition matrices for a repetition with prefactor *s*^*r*^ and the transition matrices for a random vector with prefactor 1 − *s*^*r*^.

**Fig. 6.**
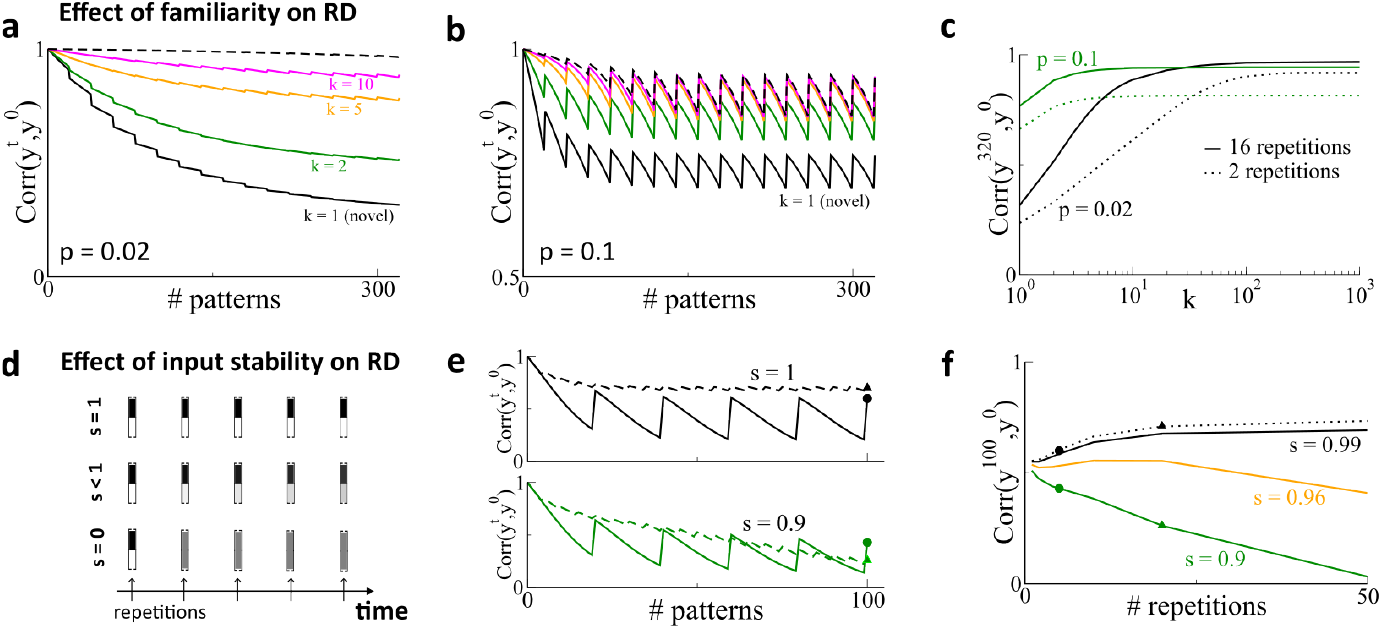
How familiarity and input stability modulate RD. **a**. We model familiarity by presenting a given pattern to the network *k* times before time *t* = 0, i.e. *k* = 1 is a novel pattern. Here *p*_+_ = *p*_−_ = 0.02, *f* = 0.15 and *N* = 1000. **b**. Same as in **a**. but with *p*_+_ = *p*_−_ = 0.1. **c**. Correlation after a total of 320 patterns, with either 16 or 2 repetitions, for low and high learning rates. **d**. The input patterns driving the observed activity may themselves undergo RD, which we parameterize with s. Here *s* = 1 indicates completely stable inputs over time while *s* = 0 indicates that the input vectors are completely uncorrelated from repetition to repetition. **e**. When inputs are stable, increased repetition rate decreases RD (*s* = 1), while this need not be the case when inputs themselves drift (*s* = 0.9). **f**. The repetition rate can reduce RD (black lines), increase RD (green line) or even leave it largely unchanged (orange line).

## Supporting information

Supplementary Figures and Detailed Calculations

## Acknowledgements

AR and FD thank Yaniv Ziv and Alon Rubin for the experimental data as well as for many helpful and enlightening discussions. AR and FD also thank Klaus Wimmer, Alex Hyafil and Jose M. Esnaola for helpful discussions and feedback. AR acknowledges “Retos” project RTI2018-097570-B-100 from the Ministry of Science and Innovation of the Spanish Government, Flag-Era project from the EU for the Human Brain Project HIPPOPLAST (Era-ICT code PCI2018-093095) and grant 2021 SGR 01522 “Dynamics in Neuronal Networks” from the Generalitat of Catalonia. This work was funded by a grant from the Spanish Ministry of Science and Innovation PID2021-124702OB-I00. This work is supported by the Spanish State Research Agency, through the Severo Ochoa and Maria de Maeztu program for Centers and Units of Excellence in R&D (CEX2020-001084-M). We thank CERCA Program/Generalitat de Catalunya for institutional support.

## Author Contributions

Conceptualization, A.R.; Methodology A.R. and F.D.; Investigation A.R., F.D., L.Z. and G.C.; Writing - Original Draft, A.R.; Writing - Review & Editing, A.R., F.D., L.Z. and G.C.; Funding Acquisition, A.R.; Resources, A.R.; Supervision, A.R.

## Additional Information

Supplementary Information File: Figures S1-S7 and detailed model description and calculations.

