## Supplementary Figures and Detailed Calculations for "Representational drift as the consequence of ongoing memory storage"

### Supplementary Information

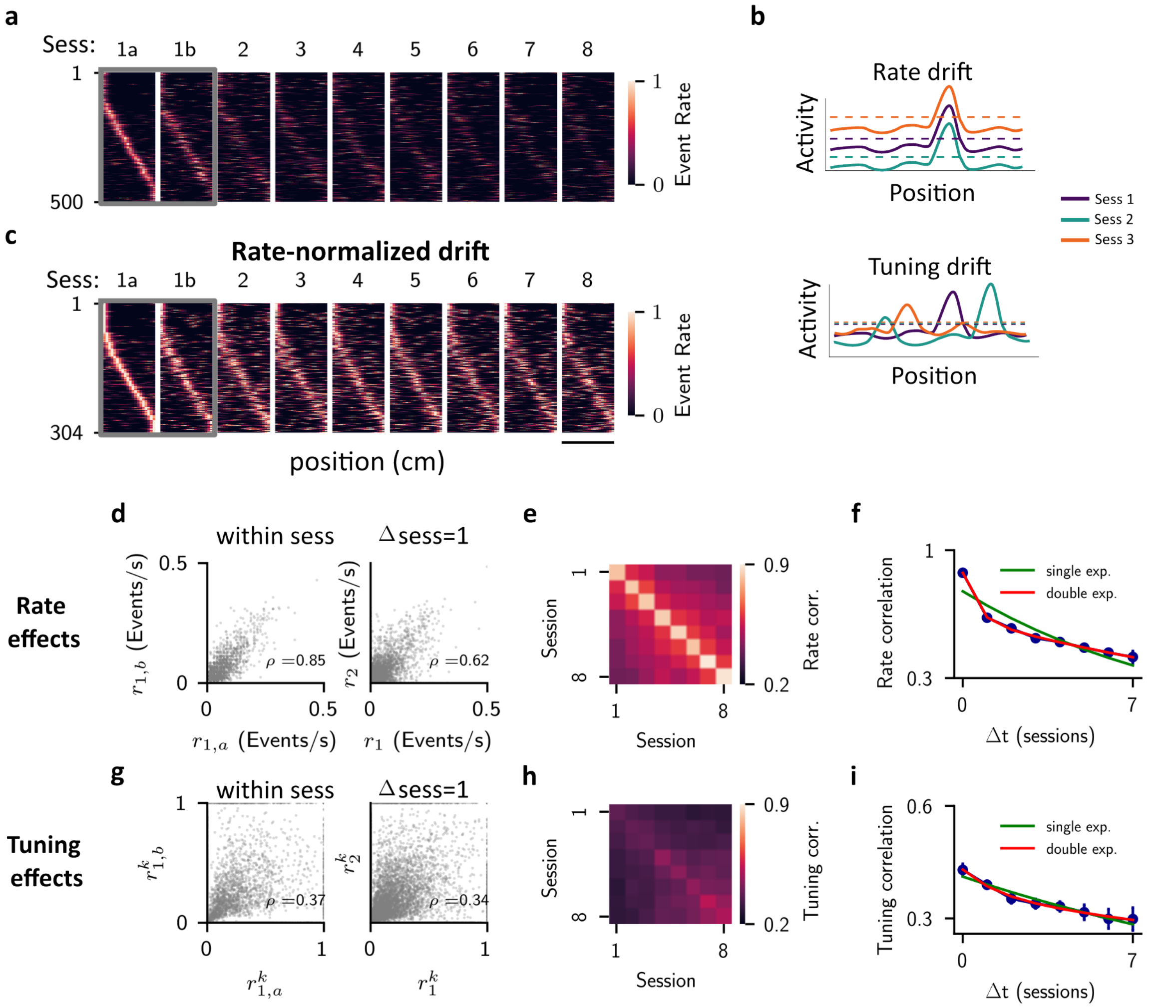


**Fig. S 1 |Analysis of data reveals rate and tuning drift occur on distinct timescales a**, Place field maps of cells active on even trials on session 1. 1a and 1b stand for even and odd trials of session 1. Cells are ordered according to their peak rate position in session 1a. Each cell event rate is normalized to the maximum reached over the 8 sessions. The scale bar is 80cm. **b**, Illustration of different changes cells may undergo over time. In the top panel, only the mean rate changes from session to session, while tuning is preserved. In the bottom panel, mean rate is preserved while tuning changes over time. **c**, Same as **a**, but plotting cells active on all sessions, and normalizing the event rate within each session. **d**, Scatter plots of mean event rates for all cells. Left: mean event rate on odd trials (ordinate axis) versus mean event rate on even trials. Right: mean event rate on session 2 (ordinate axis) versus mean event rate on session 1. $\rho$ is the Pearson correlation coefficient. **e**, Color map showing the rate correlation for all possible session pairs. The bright diagonal indicates strong within-session correlation compared to across-session correlation. **f**, Mean rate correlation as a function of elapsed sessions (blue points), averaging over all possible session pairs. A double exponential fit (red) reveals the presence of two time scales in the data. **g** Scatter plots of position-specific event rates. Left: event rate in bin $k$ on odd trials (ordinate axis) versus the event rate in the same bin on even trials. Right: event rate in bin $k$ on session 2 (ordinate axis) versus the event rate in the same bin on session 1. **h**, Same as **e**, but for tuning correlation. **i**, Same as **f**, but for tuning correlation. Note that a single time scale captures the decay over time of the correlation.

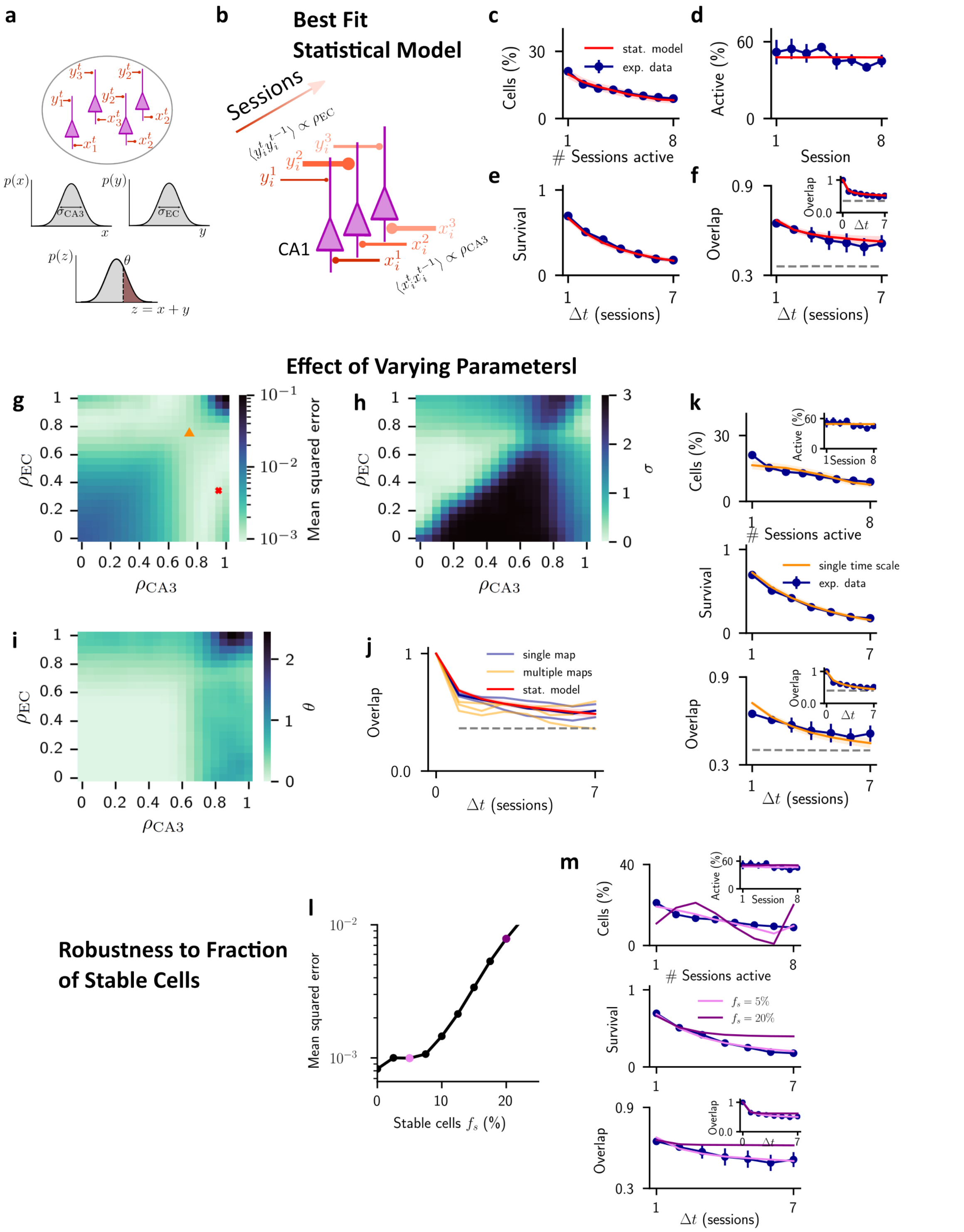


**Fig. S 2 |Modeling session-to-session changes in two sources of input with disparate temporal correlations accounts for data on active cells.** **a**, A pool of CA1 cells (in pink), where each cell $i$ receives two distinct inputs $x_i^t$ and $y_i^t$ in each session $t$. Inputs are Gaussian distributed across cells with variances $\sigma^2_{\mathrm{CA3}}$ and $\sigma^2_{\mathrm{EC}}$ respectively. Cells are active whenever their total input $z_i^t = x_i^t + y_i^t$ exceeds a threshold $\theta$. **b**,The inputs change from session to session with autocorrelations $\rho_{\mathrm{CA3}}$ and $\rho_{\mathrm{EC}}$ respectively, while preserving overall input statistics (see Methods). **c-f**, Best fit of the model (red line) to the experimental data (blue points): **c**, Distribution of the number of sessions each cell is active; **d**, Fraction of active cells over time; **e**, Survival probability; **f**, Overlap of the population activity vector. Parameters: $\theta = 0.55$, $\sigma_{CA3}/\sigma_{EC} = 1.16$, $\rho_{\mathrm{CA3}} = 0.95$, $\rho_{\mathrm{EC}} = 0.35$. **g**, Colormap showing the mean squared error of the fit in the $(\rho_{\mathrm{CA3}}, \rho_{\mathrm{EC}})$ plane. Red cross indicates parameters chosen for panels **c-f**, orange triangle parameters of panel **k**. **h,i** Colormaps showing parameters $\sigma$ and $\theta$ for the solutions found in panel **g**. **j** Overlap over time for mice with a single map per environment (blue, also shown in Fig.2) and multiple maps per environment[62] (yellow). **k**, Simulations of the statistical model where we impose both inputs to evolve on the same time scale (dark triangle in panel **a**). **l**, Error of the fit as a function of the percentage of stable cells in the model. Pink and purple dots correspond to simulations of panel **m**. **m**, Best fit of the model with 5 % (pink) and 20 % (purple) of stable cells.

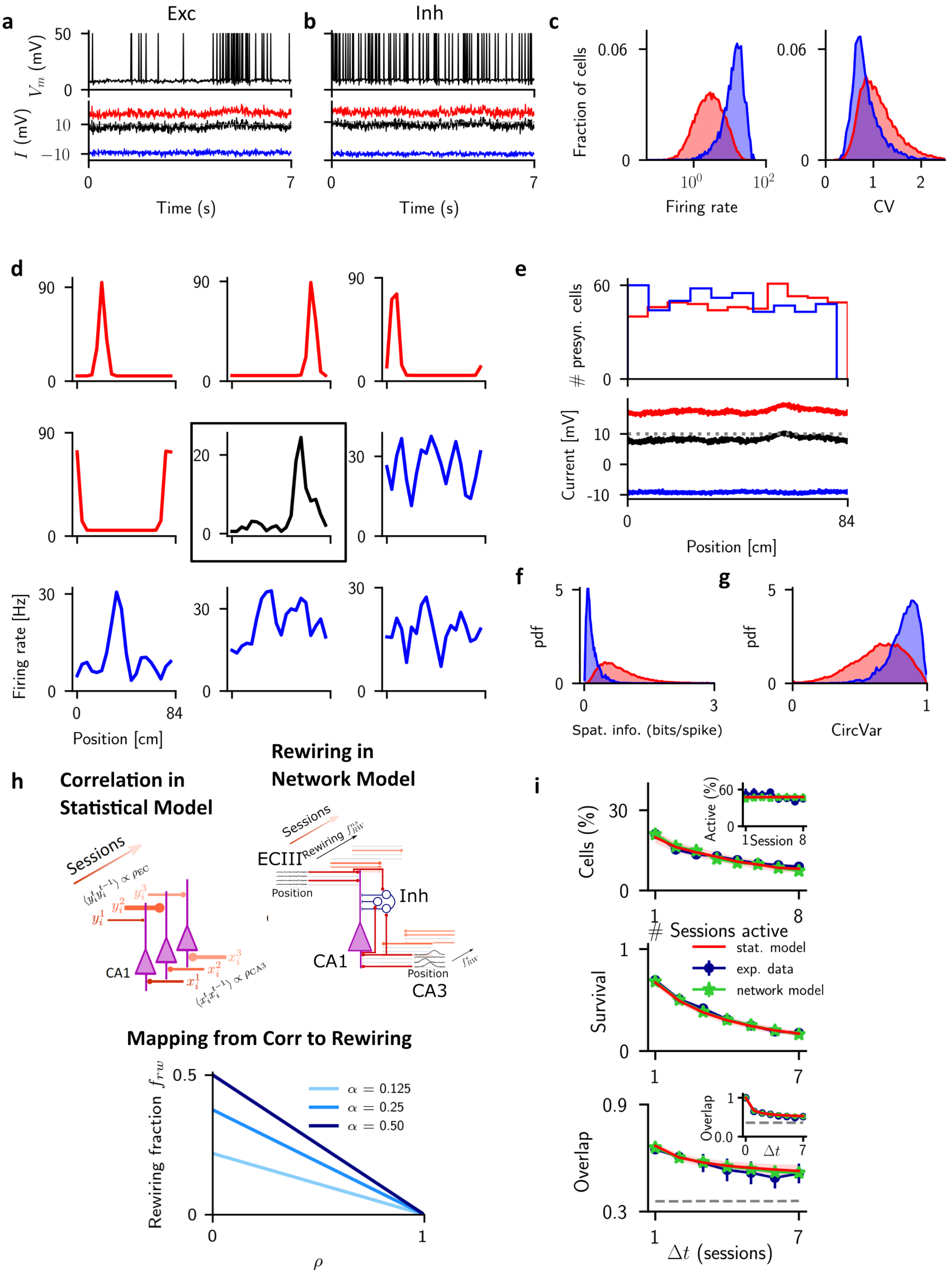

**Fig. S 3 | The network operates in the balanced regime. a,** Membrane voltage (top) and excitatory (red), inhibitory (blue) and total (black) currents (bottom) into an example excitatory neuron over one lap on the track (lap time: 7s). Excitatory and inhibitory currents balance, so that the total input is right below threshold. Shading indicated the place field of the cell. **b**, Same as **a**, but for an example inhibitory cell. **c,** Distribution of mean firing rates (left) and ISI coefficient of variations (right) for excitatory (red) and inhibitory (blue) cells. Data pooled across all sessions. Only cells with >10 spikes per session are considered. **d,** Tuning curve of the same cell as in panel **a** (black), and of 8 of its excitatory (red) and inhibitory (blue) neurons. Only tuned excitatory neurons are considered. **e,** Distribution of place field positions of all inputs to the cell highlighted in panel **d** (top), and the corresponding mean excitatory (red), inhibitory (blue) and total (black) synaptic input. **f-g** Distributions of spatial information and circular variance for all cells across the network. **h.** The relationship between the temporal correlation in the inputs in the statistical model $\rho$ and the rewiring fraction in the network model $f$. The value is given by the formula $f_{rw} = 2\alpha\,(1-\alpha)\,(1-\rho)$. **i,** The statistics for active cells, identical to those shown in Fig.S2, with the inclusion of the results from simulations of the network model.

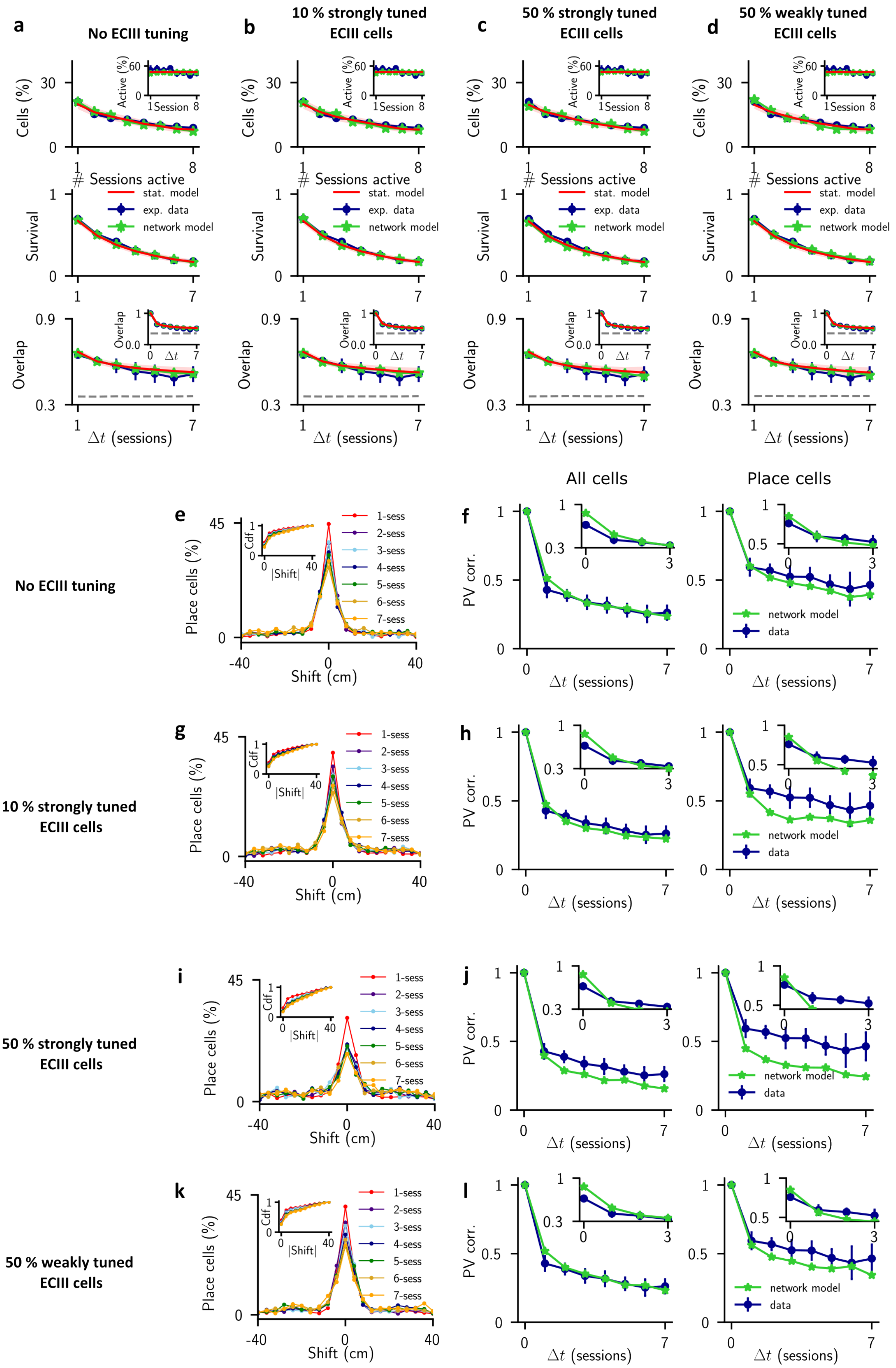
a
No ECIII tuning
b
10 % strongly tuned ECIII cells
c
50 % strongly tuned ECIII cells
d
50 % weakly tuned ECIII cells
Cells (%)
Active (%)
Session
### Sessions active
Survival
stat. model
exp. data
network model
Overlap
$\Delta t$ (sessions)
All cells
Place cells
e
No ECIII tuning
Place cells (%)
Cdf
|Shift|
Shift (cm)
1-sess
2-sess
3-sess
4-sess
5-sess
6-sess
7-sess
f
PV corr.
network model
data
g
10 % strongly tuned ECIII cells
h
i
50 % strongly tuned ECIII cells
j
k
50 % weakly tuned ECIII cells
l

**Fig. S 4 | The effect of tuning in EC on RD. a,** Adding tuning to EC does not alter the statistics of active cells. **b,**: Increasing the tuning in EC cells leads to increased diffusion in the position of place cell centroids (decreased peak in histogram of centroids shifts), and a sharper drop in PV correlation of place cells. This affect is most prominent when cells in EC are tuned according to a von Mises distribution, as in CA3, i.e. "strongly tuned". Weaker tuning, of a cosine type, does not lead to significant effects, even if the fraction of tuned cells is large, see bottom row. All parameters are as in Fig.3.

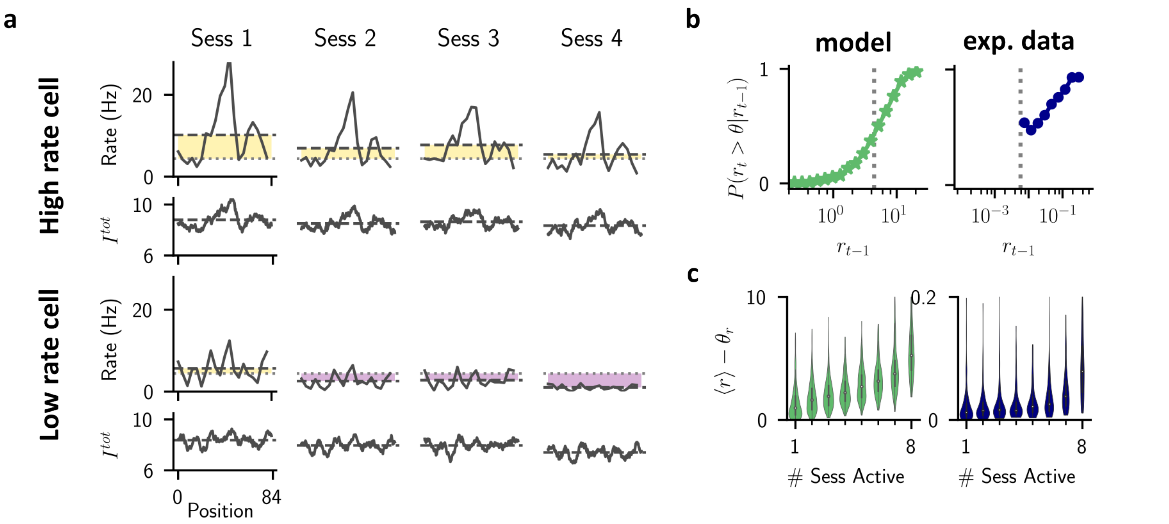


**Fig. S 5 | The correlation between firing rate and stability predicted in network model is observed in the data a.** The firing rate and mean total input for two sample cells with high (top) and low (bottom) mean firing rate. Dashed lines indicate averages over all spatial positions, while the light gray dotted line in the threshold. Yellow and purple shadings highlight the distance from threshold, from above and below respectively. **b.** The probability of a cell being above threshold (for detection as active) conditioned on its mean firing rate from the previous session, in the network model (left) and experimental data (right). **c.** Distributions of mean firing rates across sessions, for cells active on different numbers of sessions, in the network model (left) and in the data (right).

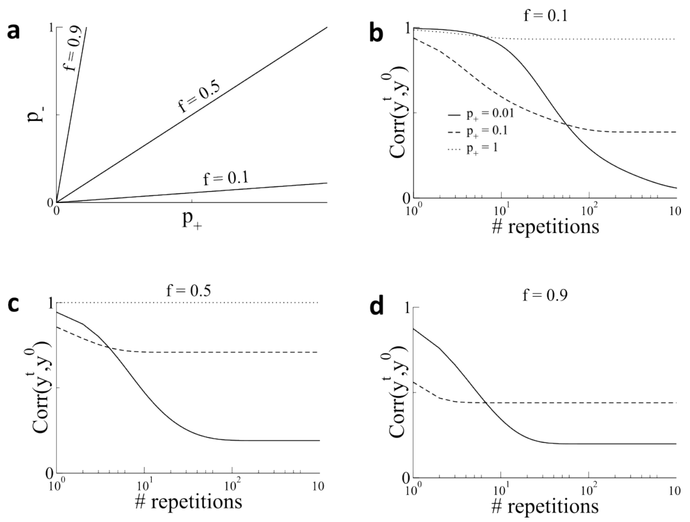


**Fig. S 6 | The effect of coding sparseness and learning rates on RD a.** The parameters are varied so as to keep the mean connectivity $\bar{c}$ fixed. **b.** Low coding sparseness, $f = 0.1$. **c.** Cofing sparseness $f = 0.5$. For this value the learning rates for potentiation and depression are the same, i.e. $p_- = p_+$. **d.** Coding sparseness $f = 0.9$.

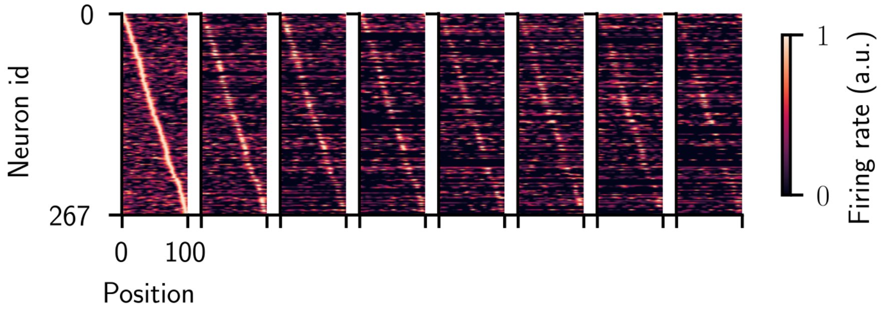


**Fig. S 7 | RD in a network model with Hebbian Plasticity** Snapshots of the activity of place cells in a network of threshold linear neurons with Hebbian plasticity. Parameters are chosen to match the decrease in the degree correlation $\rho_d$ from the spiking network, see Methods and Fig.3.

### Detailed model description and calculations

We characterized how spatially tuned versus untuned features of neuronal activity differentially contributed to the observed drift in the data. In the case of mice running on a linear track, using the ordering of place cells from the first session shows that there is significant drift on subsequent sessions[7], Fig.2a.

We asked to what extent this drift was due to shifts in the place-field location of cells or simply to changes in rate, Fig.2b. To do so we normalized the time-averaged spatial profile of each place cell by its maximal firing rate on each session. Doing so revealed much less drift, and specifically there was no longer a large difference between the drift from session 1 to session 2 and the degree of variability within session 1, Fig.2c. To quantify this effect we compared scatter plots of the mean event rate of all cells within session 1, and then between sessions 1 and 2, Fig.2d. The correlation of the mean rates evaluated on only even trials versus only odd trials within the session 1 was much higher than the correlation of mean rates between sessions 1 and 2. The pronounced drop in correlation occurred reliably from one session to the next regardless of the session identity, as can be seen in the matrix of rate correlations in Fig.2e. The correlation as a function of session separation averaged over all sessions showed a steep initial drop, followed by a slower decay, Fig.2f. In fact, the curve was much better fit by a double exponential than a single exponential suggesting the presence of two processes operating at distinct time-scales.

On the other hand, when we compared mean rates for each spatial bin of the rate-normalized profiles, we found a much smaller decrease in correlation, Fig.2g. Again this drop was slight from one session to the next irrespective of the session identity, and the curve of the average correlation as a function of session separation showed a shallow decay which was much better fit by a single exponential function than the non-normalized curve, Fig.2h-i. The time-scale of this decay was the same as that of the slower of the two time-scales of the mean rates in Fig.2f. This strongly suggested that the slow time-scale decay in the overall RD was due to changes in the spatial tuning of cells, while the fast drop-off in correlation from one session to the next was related to changes in mean activity.

#### Statistical Model

We hypothesized that these two time-scales might reflect plasticity along the two main afferent excitatory pathways to CA1: Schaffer collaterals from CA3, and inputs originating from layer III of entorhinal cortex (EC). To test this we investigated if changes in synaptic input alone could account for key experimental findings in a simple, statistical model. Specifically, we considered a population of binary neurons, each one of which received two inputs on any given session $t$, which we call $x^t$ and $y^t$, Fig.2a. On the first session we distributed these inputs randomly across neurons according to Gaussian distributions with zero mean and variances $\sigma^2_{CA3}$ and $\sigma^2_{EC}$ respectively. This led naturally to a large degree of heterogeneity in the inputs to single cells, with some receiving strong excitatory drive and others being strongly inhibited. The total input to a cell $z^t = x^t + y^t$ was then compared to a threshold $\theta$ to determine if the cell was active or not, i.e. cell activity was binary. Therefore, in this simplified model we did not take continuous changes in the event rate of the cells into account, nor did we explicitly include spatial tuning. To model RD we drew these inputs anew from session to session from the same Gaussian distributions, but allowed for some temporal correlation. Namely, the session-to-session autocorrelation for inputs from CA3 was

$$\rho_{CA3} = \langle x^t x^{t-1} \rangle / \sigma^2_{CA3}, \tag{16}$$

where the brackets indicate an average over all neurons, Fig.2b, and there is an equivalent relation for inputs from EC. There were four model parameters in total after normalization: the ratio of the variances, the threshold, and the two autocorrelations. We optimized these parameters to fit four characteristic measures of neuronal activity from the experimental data: 1 - the distribution of the number of sessions in which a cell is active, 2 - the fraction of active neurons per session, 3 - Survival fraction. This measure includes only those cells which are initially active in the first session, and quantifies the fraction of these which continue to be active on subsequent sessions, and 4 - the overlap in the vector of active neurons from session to session. The model was able to reproduce these measures quantitatively Fig.2c-f, but only when the temporal correlation along one input pathway was close to one ($\rho = 0.95$) and the other sufficiently small ($\rho = 0.35$), Fig.S2g-i. In this regime, one of the two inputs changed only very slowly from session to session while the other was almost entirely redrawn randomly for each session, consistent with the two disparate time-scales we observed from direct analysis of the data. Because the model only considered mean inputs, and cell activity was just the sum of the two, it was not possible to say which of the two inputs should vary slowly over time, and which quickly. Once spatial tuning was taken into account (see next section), fast plasticity in EC inputs and slow plasticity in CA3 inputs was the only choice compatible with the data. Finally, we note that a small fraction of cells in the data were active on all 8 sessions, Fig.2c, raising the possibility of a sub-population of stable cells which reliably participated in the hippocampal code. We modeled this by assuming that a fraction $f_s$ of cells were always active, and then allowing the remaining fraction $1 - f_s$ to evolve according to the model described above. We found that the mean squared error of the model fit simultaneously to the four above-mentioned measures increased for increasing $f_s$, but that it remained low when the stable fraction was below 10 percent, Fig.S2l. Therefore, while the model fit was always best without dedicated stable cells, the data would nonetheless also be consistent with a small fraction of stable cells.

#### Input dynamics as an Ornstein-Uhlenbeck process

The update rules Eq.1 can be rewritten, in continous time, in this form:

$$\epsilon \frac{dz}{dt} = \sqrt{1-\rho^2}\sigma\eta(t) - (1-\rho)\, z(t), \tag{17}$$

where $\langle \eta(t) \rangle = 0$ and $\langle \eta(t)\eta(s) \rangle = \delta(t-s)$.
Eq. (17) is a Ornstein-Uhlenbeck with drift coefficient $\mu = (1-\rho)/\epsilon$ and diffusion coefficient $D = \sigma^2 \left(1-\rho^2\right)/(2\epsilon^2)$. The associated Fokker-Planck equation for the probability distribution of the input $P(z,t)$ is then:

$$\frac{\partial P(z,t)}{\partial t} = \mu \frac{\partial\,(zP)}{\partial z} + D\frac{\partial^2 P}{\partial z^2}. \tag{18}$$

The stationary solution of equation (18) is a gaussian distribution centered at zero with variance

$$\sigma^2 = \frac{D}{\mu} = \frac{\sigma^2\left(1-\rho^2\right)}{2\epsilon\,(1-\rho)}. \tag{19}$$

We therefore set $\epsilon = \left(1-\rho^2\right)\left[2\,(1-\rho)\right]^{-1}$ so that, if the initial distribution of $z$ is $\mathcal{N}(0,\sigma)$ the process is stationary. Eq. (17) is then

$$\frac{dz}{dt} = \sqrt{2D}\eta(t) - \mu z(t), \tag{20}$$

where

$$\mu = \frac{2}{(1-\rho^2)}\left[1-\rho\right]^2 \quad D = \sigma^2\mu. \tag{21}$$

In the limit $p \to 0$, we have $dz/dt = 0$. When $p$ goes to 1, $dz/dt = 2(\sigma\eta - z)$.
The covariance of a process of the type Eq.(17) is given by:

$$\langle z(t)z(0) \rangle = \sigma^2 e^{-t/\tau}, \tag{22}$$

where we defined $\tau = \mu^{-1}$. Note that $\lim_{\rho\to 0}\tau = 1/2$ and $\lim_{\rho\to 1}\tau = \infty$: when there is no plasticity, the autocorrelation decay time is infinite.

#### Network equations

Each model neuron follows the LIF dynamics:

$$\tau\dot{V}_i^l = -V_i^l + \tau I_i^l \tag{23}$$

with $i = 1, \dots, N_l$ and $l \in \{E, I\}$. $I_i^l$ is the synaptic current. The neuron emits a spike when it crosses $V_{\text{th}} = 10$ mV, where it is then resetted to $V_r = 0$ mV.
The synaptic current to excitatory neurons is:

$$I_i^E = \frac{g_s}{\tau_s} \sum_{j=1}^{N_{\text{CA3}}} c_{ij}^{E,s} \sum_k e^{-\left(t - t_{j,k}^s\right)/\tau_s} - \frac{g_{EI}}{\tau_s} \sum_{j=1}^{N_I} c_{ij}^{E,I} \sum_k e^{-\left(t - t_{j,k}^I\right)/\tau_s} + \frac{g_{ns}}{\tau_s} \sum_{j=1}^{N_{\text{EC}}} c_{ij}^{E,ns} \sum_k e^{-\left(t - t_{j,k}^{ns}\right)/\tau_s} \tag{24}$$

where $g_{s,EI}$ are the feedforward and E $\leftarrow$ I synaptic conductances, $t_{j,k}^l$ are the times of the $k$th action potential of neuron $j$ of the $l$ population, and $C^{l,r}$ the connectivity matrices.
The synaptic current to inhibitory neurons is:

$$I_i^I = \frac{g_{IE}}{\tau_s} \sum_{j=1}^{N_E} c_{ij}^{I,E} \sum_k e^{-\left(t - t_{j,k}^E\right)/\tau_s} - \frac{g_{II}}{\tau_s} \sum_{j=1}^{N_I} c_{ij}^{I,I} \sum_k e^{-\left(t - t_{j,k}^I\right)/\tau_s} + \frac{g_{Is}}{\tau_s} \sum_{j=1}^{N_{\text{CA3}}} c_{ij}^{E,s} \sum_k e^{-\left(t - t_{j,k}^s\right)/\tau_s} + \frac{g_{Ins}}{\tau_s} \sum_{j=1}^{N_{\text{EC}}} c_{ij}^{E,ns} \sum_k e^{-\left(t - t_{j,k}^{ns}\right)/\tau_s} \tag{25}$$

Unless otherwise stated, we set $g_{Is} = g_s$ and $g_{Ins} = g_{ns}$. To set the network in the so called balance state, the synaptic weights are scaled according to:

$$g_l = \frac{G_l}{\sqrt{K}}, \tag{26}$$

where $K$ is the average number of active inputs to each subpopulation, and $G_l$ is independent of $K$

#### Network parameters

The membrane time constant is set to $\tau = 10$ ms, and the synaptic time constant $\tau_s = 5$ ms. The synaptic weights of CA1 recurrent connections are $G_{EI} = 2.8$ mV, $G_{IE} = 2.0$ mV, $G_{II} = 2.0$ mV[1]. The other synaptic weights (that determine the variances of spatial and non-spatial inputs) are set in the following considering the fit of the statistical model. Parameters of the CA3 place fields are $\beta = 19.85$, $r_b = 5$ Hz, $\bar{R} = 90$ Hz. Synaptic weights were defined as $g_{CA3} = \frac{G_{CA3}}{\sqrt{f\alpha N}}$ and $g_{EC} = \frac{G_{EC}}{\sqrt{f\alpha N}}$, where $G_{CA3} = 3.5$ and $G_{EC} = G_{CA3}/1.16$.
Unless specified otherwise, we consider $N_E = 4000$ and $N_I = 1000$ CA1 cells. The connections probability is the same for all connectivity matrices and equal to $\alpha = 0.125$. The sparsity of the CA3 and EC layer are set respectively to $f_{CA3} = 0.5$ and $f_{EC} = 0.5$. We wish to have about the same number of excitatory CA1, CA3 and EC active cells, so that we set $N_{CA3} = N_E/f_{CA3} = 8000$ and $N_{EC} = N_E/f_{EC} = 8000$. The fraction of tuned CA3 inputs $f_s$ is set to $f_s = 0.5$, unless specified otherwise.

#### Input correlations in the network model

In this appendix, we calculate the amount of correlations of the inputs expected in the network model, for the same or different environments. In the network model, average synaptic inputs to a cell $i$ in environment $A$ can be written in the form:

$$I_i^A = \sum_j c_{ij}^A \nu_j^A, \tag{27}$$

[1]Note that these weights are then rescaled according to Eq.26.

where we omitted conductances and time constants. $c_{ij}^A$ is the matrix element during visits of environment $A$ (if visits do not occur at the same time plasticity may have occurred), and $\nu_j^A$ is the firing rate of input neuron $j$ in environment $A$.

In general, the Pearson correlation of inputs from different environments can be written as:

$$\rho = \frac{\langle I^A I^B \rangle - \mu_A \mu_B}{\sigma_A \sigma_B}, \tag{28}$$

where we have defined $\mu_{A,B} = \langle I^{A,B} \rangle$ and dropped the index $i$ for simplicity. Note that the bracket averages are all over the index $i$ (i.e. we consider one specific realization of the input neurons rate $\nu$). The expected values take the form:

$$\mu_{A,B} = \sum_j \langle c_j^{A,B} \nu_j^{A,B} \rangle = \alpha \sum_j \nu_j^{A,B}, \tag{29}$$

where for the second equality we have assumend that the connection probability $\alpha$ in the two environments is the same. On the other hand, we have for the expected value of the product of the inputs:

$$\langle I^A I^B \rangle = \sum_j \sum_l \langle c_j^A c_l^B \nu_j^A \nu_l^B \rangle = \sum_j \langle c_j^A c_j^B \rangle \nu_j^A \nu_j^B + \sum_{j \neq l} \langle c_j^A c_l^B \rangle \nu_j^A \nu_l^B. \tag{30}$$

We then have

$$\langle I^A I^B \rangle = \sum_j \nu_j^A \nu_j^B \langle c_j^A c_j^B \rangle + \alpha^2 \sum_{j \neq l} \nu_j^A \nu_j^B. \tag{31}$$

For the within-environment variances, we have:

$$\left(\sigma^{A,B}\right)^2 = \sum_j \sum_l \nu_j \nu_l \langle c_j c_l \rangle - \mu^2 = \alpha \sum_j \nu_j^2 + \alpha^2 \sum_{j \neq l} \nu_j \nu_l - \alpha^2 \sum_{j,l} \nu_j \nu_l, \tag{32}$$

where we dropped the indeces A,B for simplicity and substituted the previously found expression for $\mu$. Adding and removing the element where $j = l$ in the second term gives:

$$\sigma^2 = \alpha \left(1 - \alpha\right) \sum_j \nu_j^2. \tag{33}$$

Using the same trick in the numerator of adding and removing one element from the sum, we get for the correlation:

$$\rho = \frac{\sum_j \nu_j^A \nu_j^B \left(\langle c_j^A c_j^B \rangle - \alpha^2\right)}{\alpha \left(1 - \alpha\right) \left[\sum_j \left(\nu_j^A\right)^2\right]^{1/2} \left[\sum_j \left(\nu_j^B\right)^2\right]^{1/2}}. \tag{34}$$

If we now consider that the input layer is very large, then all sums over $j$ can be rewritten in terms of mean values. If we further assume that the input patterns are drawn from the same distribution for the different environments, then we have:

$$\rho = \frac{\langle \nu_j^A \nu_j^B \rangle_j \left[\langle c_j^A c_j^B \rangle - \alpha^2\right]}{\langle \nu_j^2 \rangle_j \alpha \left(1 - \alpha\right)}, \tag{35}$$

where $\langle \cdot \rangle_j$ stands for the average over the input layer.

Eq.(35) is the general form of the inputs considering different environments/directions or the same environment at different times. We now consider three separate cases:

1. Visits of the same environment at different times (same inputs firing rates but different connectivity matrices)

2. Partially correlated environments at approximately the same time (different input firing rates but constant connectivity)

3. Orthogonal environment at different times.

**Case 1.** If the environment is the same, at least for the spatial input we can consider the input firing rates to be constant over visits (in first approximation). This implies $\langle \nu_j^A \nu_j^B \rangle_j = \langle \nu_j^2 \rangle_j$, which then gives:

$$\rho_t = \frac{\langle c_j^t c_j^{t+1} \rangle - \alpha^2}{\alpha\,(1-\alpha)}, \tag{36}$$

where $t$ stands for the time of the visit. Eq. (36) can be made more explicit by expressing $\langle c_j^t c_j^{t+1} \rangle$ in terms of probabilities. In fact, since $c_j$'s are binary matrix elements, we can write:

$$\langle c_j^t c_j^{t+1} \rangle = \text{Pr}\left(c_j^t = 1, c_j^{t+1} = 1\right) = \text{Pr}\left(c_j^{t+1} = 1 \middle| c_j^t = 1\right) \text{Pr}\left(c_j^t = 1\right). \tag{37}$$

Since we have $\text{Pr}\left(c_j^t = 1\right) = \alpha$, we then have

$$\rho_t = \frac{\text{Pr}\left(c_j^{t+1} = 1 \middle| c_j^t = 1\right) - \alpha}{1-\alpha}. \tag{38}$$

If there is no plasticity, then $\text{Pr}\left(c_j^{t+1} = 1 \middle| c_j^t = 1\right) = 1$ and $\rho_t = 1$, while if there is a complete rewiring $\text{Pr}\left(c_j^{t+1} = 1 \middle| c_j^t = 1\right) = \alpha$ and $\rho_t = 0$.

**Case 2.** In this case, we assume that the connectivity matrix is the same for the different visits, and the input patterns are partially correlated. In the data we have, this would for example be the case of considering the two running directions as different environments. We then have here that $\langle c_j^A c_j^B \rangle = \alpha$, so that the correlation is

$$\rho_{LR} = \frac{\langle \nu_j^L \nu_j^R \rangle_j}{\langle \nu_j^2 \rangle_j}, \tag{39}$$

where $L$ and $R$ stands for left and right. Input neurons $j$ are either silent or active with firing rate $\nu$, so that we have $\langle \nu_j^2 \rangle_j = f\nu^2$, where $f$ is the sparsity of the input layer (fraction of active cells). As for the previous case, we can write more explicit the correlation using probabilities:

$$\rho_{LR} = \frac{\nu^2 \text{Pr}\left(\nu^L = \nu, \nu^R = \nu\right)}{f\nu^2} = \text{Pr}\left(\nu^L = \nu \middle| \nu^R = \nu\right); \tag{40}$$

the correlation between running directions is just the probability that given an input cell is active in one running idrection, it is active also in the other.

**Case 3**. Since environments are assumed to be orthogonal or independent, then we have $\langle \nu^A \nu^B \rangle = f\nu^2$, so that we get for the correlation

$$\rho_{AB} = \frac{f\left(\langle c^A c^B \rangle - \alpha^2\right)}{\alpha\,(1-\alpha)}. \tag{41}$$

We summarize here the three correlations found:

- $\rho_t = \dfrac{\text{Pr}\left(c_j^{t+1} = 1 \middle| c_j^t = 1\right) - \alpha}{1-\alpha}$,
- $\rho_{LR} = \text{Pr}\left(\nu^L = \nu \middle| \nu^R = \nu\right)$,
- $\rho_{AB} = \dfrac{f\left(\langle c^A c^B \rangle - \alpha^2\right)}{\alpha\,(1-\alpha)}$.

#### Steady-state Connectivity Statistics in Hebbian plasticity model

We assume that the learning process has gone on long enough so that the connectivity has reached a stationary state and hence is independent of the initial configuration. Then we consider the encoding of one pattern and calculate the mean and the variance in the number of inputs to a cell in the postsynaptic layer (CA1). We assume $N$ neurons in the presynaptic layer, e.g. EC, and patterns with sparseness $f_{\rm pre} = f_{\rm post} = f$ for simplicity. The synapse between two co-active cells is potentiated with probability $p_+$, and a synapse between one active and one inactive cell is depressed with probability $p_-$. Synapses between inactive cells are not updated. We consider binary synapses such that the connection from cell $j$ in the presynaptic layer to cell $i$ in the postsynaptic layer is $c_{ij} \in \{0, 1\}$. The degree of a cell $i$ can be written $d_i = \sum_{j=1}^{N} c_{ij}$. The mean degree in the network is then $\mu_d = \langle d_i \rangle = N\langle c_{ij} \rangle = N\bar{c}$, and the average over synapses can be written as

$$\bar{c} = f_{\rm post}\langle c_{ij} \rangle_{i\in\Omega_f} + (1 - f_{\rm post})\langle c_{ij} \rangle_{i\notin\Omega_f}, \tag{42}$$

where $\omega_f$ is the set of neurons which are part of the current, activated pattern. By considering the probability that a synapse is in the potentiated state depending on whether or not the neuron is part of the pattern, one arrives at

$$\langle c_{ij} \rangle = \frac{f^2 p_+}{f^2 p_+ + 2f(1-f)p_-} \tag{43}$$

The variance in the degree $\sigma_d^2 = \langle d_i^2 \rangle - \mu_d^2$. The first term can be written

$$\langle d_i^2 \rangle = \langle \sum_j c_{ij}^2 + \sum_j \sum_{k\neq j} c_{ij}c_{ik} \rangle = N\langle c_{ij}^2 \rangle + N(N-1)\langle c_{ij}c_{ik} \rangle$$

The first term $\langle c_{ij}^2 \rangle = \langle c_{ij} \rangle = \bar{c}$, while the second term can be calculated by considering the probability of both connections being potentiated as a function of whether or not the pre- and postsynaptic cells are active in the current pattern, or not. For simplicity we write $\bar{c}_{11} = \langle c_{ij}c_{ik} \rangle = Pr(c_{ij} = 1, c_{ik} = 1)$. The result is

$$\bar{c}_{11} = \frac{f^2 p_+\Big(f(p_+ + 2(1-p_+)\bar{c}) + 2(1-f)(1-p_-)\bar{c}\Big)}{1 - f\big(f(1-p_+) + (1-f)(1-p_-)\big)^2 - (1-f)(1-fp_-)^2} \tag{44}$$

Finally, the autocorrelation of the degree can be written

$$\rho_d = \frac{\langle d_i^t d_i^{t-1} \rangle - \mu_d^2}{\sigma_d^2}, \tag{45}$$

where we can write $\langle d_i^t d_i^{t-1} \rangle = N\langle c_{ij}^t c_{ij}^{t-1} \rangle + N(N-1)\langle c_{ij}^t c_{ik}^{t-1} \rangle$. The two covariances can, once again, be calculated by considering the probability that the product is equal to one depending on the participation of the cells in the patterns at time $t-1$ and $t$. The result is

$$\langle c_{ij}^t c_{ij}^{t-1} \rangle = \bar{c}\Big(1 - 2p_- f(1-f)\Big), \tag{46}$$

$$\langle c_{ij}^t c_{ik}^{t-1} \rangle = \bar{c}f^2 p_+ + \bar{c}_{11}\Big(1 - f^2 p_+ - 2p_- f(1-f)\Big) \tag{47}$$

#### Analytical solution for RD in Plasticity Model

The relative simplicity of the plasticity model allows us to calculate how an output pattern changes in time, given a particular input. The drop in correlation of the output pattern, or the drift in the representation, can therefore be calculated analytically. We recall that the parameters of the model are the learning rates $p_+$ and $p_-$, the coding fraction $f$, and the population size $N$. The theoretical solution will be exact in the limit $N \to \infty$. We will consider the scenario for which the network has reached a steady state after many statistically equivalent patterns have been encoded. We then encode the pattern to be tracked, and allow for an arbitrary protocol of repetitions and interleaved random patterns. In this way the effects of familiarization, repetition rate as well as the number of interleaved unrelated episodes can all be studied.

We calculate the correlation between the activity of the network at time $t$ and the activity at time $t = 0$ which can be written

$$\mathrm{Corr}(y^t, y^0) = \frac{\mathrm{Cov}(y^t, y^0)}{\sqrt{\mathrm{Var}(y^t)\mathrm{Var}(y^0)}}, \tag{48}$$

where

$$\mathrm{Cov}(y^t, y^0) = \frac{1}{N}\sum_{i=1}^{N} y_i^t y_i^0 - \bar{y}^t \bar{y}^0, \tag{49}$$

and $\mathrm{Var}(y^t) = \mathrm{Cov}(y^t, y^t)$, while the mean output is

$$\bar{y}^t = \frac{1}{N}\sum_{i=1}^{N} y_i^t. \tag{50}$$

The vector $y^t$ is the output vector of the network at time $t$ in response to an input vector which we call $x \in \{0, 1\}$. For simplicity we assume that the input vector does not change, and hence the activity of a cell $i$ is

$$y_i^t = \frac{1}{N}\sum_{j=1}^{N} c_{ij}^t x_j, \tag{51}$$

where $c_{ij}^t = 1$ if there is a synaptic connection, or else $c_{ij}^t = 0$. We can choose the ordering of the presynaptic and postsynaptic cells such that pattern we are tracking is the one where the first $fN$ cells are the active ones. In that case

$$y_i^t = \frac{1}{N}\sum_{j=1}^{fN} c_{ij}^t, \tag{52}$$

where the mean and variance of the input will depend on whether $i \leq fN$ or not. For $i \leq fN$ the cell will have been active in the tracked memory. Specifically, when the tracked memory is encoded, we have ordered the cells so that in the first quadrant of the connectivity matrix there is an $fN$ by $fN$ size block in which the synapses have been subjected to potentiations. Below this there is an $(1 - f)N$ by $fN$ block where the synapses have been subjected to depressions. The remaining two blocks in the matrix are not relevant for determining the output correlation since the input $x$ is a vector of ones for the first $fN$ entries and otherwise zero.

The mean input $\bar{y}^t$ is

$$\begin{aligned} \bar{y}^t &= \frac{1}{N}\sum_{i=1}^{N} y_i^t, \\ &= \frac{1}{N}\sum_{i=1}^{fN} y_i^t + \frac{1}{N}\sum_{i=fN+1}^{N} y_i^t, \\ &= f\langle y^t\rangle_P + (1 - f)\langle y^t\rangle_D, \\ &= f^2\langle c^t\rangle_P + f(1 - f)\langle c^t\rangle_D, \end{aligned} \tag{53}$$

where for the last line we have used Eq.52, and the subscripts $P$ and $D$ refer to the statistics of inputs which have been potentiated or depressed during the storage of the tracked memory, respectively.

The variance of the input is

$$\begin{aligned}
\mathrm{Var}(y^t) &= \frac{1}{N}\sum_i (y_i^t)^2 - (\bar{y}^t)^2, \\
&= \frac{1}{N}\sum_{i=1}^{fN}(y_i^t)^2 + \frac{1}{N}\sum_{i=fN+1}^{N}(y_i^t)^2 - (\bar{y}^t)^2, \\
&= \frac{f}{N^2}\Big(\sum_{j=1}^{fN}\langle c_{ij}^t\rangle_P + \sum_{j=1}^{fN}\sum_{k\neq j}\langle c_{ij}^t c_{ik}^t\rangle_P\Big) + \frac{1-f}{N^2}\Big(\sum_{j=1}^{fN}\langle c_{ij}^t\rangle_D + \sum_{j=1}^{fN}\sum_{k\neq j}\langle c_{ij}^t c_{ik}^t\rangle_D\Big) - (\bar{y}^t)^2, \\
&= \frac{f^2}{N}\Big(\langle c^t\rangle_P - \langle cc^t\rangle_P\Big) + f^3\langle cc^t\rangle_P + \frac{f(1-f)}{N}\Big(\langle c^t\rangle_D - \langle cc^t\rangle_D\Big) + f^2(1-f)\langle cc^t\rangle_D \\
&\quad -(\bar{y}^t)^2,
\end{aligned} \tag{54}$$

where for simplicity we have written $\langle c_{ij}^t c_{ik}^t\rangle = \langle cc^t\rangle$.

The covariance of the input at times 0 and $t$ is

$$\begin{aligned}
\mathrm{Cov}(y_t, y_0) &= \frac{f^2}{N}\Big(\langle c_{ij}^t c_{ij}^0\rangle_P - \langle c_{ij}^t c_{ik}^0\rangle_P\Big) + f^3\langle c_{ij}^t c_{ik}^0\rangle_P + \frac{f(1-f)}{N}\Big(\langle c_{ij}^t c_{ij}^0\rangle_D - \langle c_{ij}^t c_{ik}^0\rangle_D\Big) \\
&\quad + f^2(1-f)\langle c_{ij}^t c_{ik}^0\rangle_D - \bar{y}^t\bar{y}^0,
\end{aligned} \tag{55}$$

#### Markov Processes for 1st and 2nd-order Statistics of Synapses

*i. Mean weight in potentiated quadrant,* $\langle c^t\rangle_P$:

A single presentation of the tracked pattern will shift the steady-state probabilities in the potentiated quadrant according to

$$\mathbf{c_P} = \mathbf{R_P}\mathbf{c_{ss}}, \tag{56}$$

where $\mathbf{c_{ss}} = (\bar{c}, 1-\bar{c})$ and

$$\mathbf{R_P}\begin{pmatrix} 1 & p_+ \\ 0 & 1-p_+ \end{pmatrix}. \tag{57}$$

The mean weight at time $t$ is then given by

$$\langle c^t\rangle = \mathbf{e_1}^{\mathrm{T}}\mathbf{A}^t\mathbf{c_P}, \tag{58}$$

where

$$\mathbf{A} = \begin{pmatrix} 1-2f(1-f)(1-p_-) & f^2p_+ \\ 2f(1-f)p_- & 1-f^2p_+ \end{pmatrix}, \tag{59}$$

and $\mathbf{e_1} = (1,0)$.

*ii. Mean weight in depressed quadrant,* $\langle c^t\rangle_D$:

The Markov process is identical, with the sole difference being the initial condition, where here $\mathbf{c_D} = \mathbf{R_D}\mathbf{c_{ss}}$ with

$$\mathbf{R_D}\begin{pmatrix} 1-p_- & 0 \\ p_- & 1 \end{pmatrix}. \tag{60}$$

*iii. 2nd-order statistic for weights in potentiated quadrant,* $\langle cc^t\rangle_P$

A single presentation of the tracked pattern shift the steady-state 2nd-order statistics in the potentiated quadrant according to

$$\xi_{\mathbf{P}} = \mathbf{Q_P}\xi_{\mathbf{ss}}, \tag{61}$$

with $\xi_{\mathbf{ss}} = (\bar{c}_{11}, \bar{c}_{01}, \bar{c}_{00})$, where $\bar{c}_{11}$ is given by Eq.44,

$$\bar{c}_{00} = \frac{f(1-f)\Big(p_-^2 + 2p_-(1-\bar{c})(2-p_- - fp_+)\Big)}{1 - f(1-fp_+ - (1-f)p_-)^2 - (1-f)(1-fp_-)^2} \tag{62}$$

and $\bar{c}_{01} = (1 - \bar{c}_{11} - \bar{c}_{00})/2$. The transition matrix for a repetition is

$$\mathbf{Q_P} = \begin{pmatrix} 1 & 2p_+ & p_+^2 \\ 0 & 1 - p_+ & p_+(1 - p_+) \\ 0 & 0 & (1 - p_+)^2 \end{pmatrix}. \tag{63}$$

The 2nd-order statistic at time $t$ is then given by

$$\langle cc^t \rangle_P = \mathbf{e_1}^{\mathrm{T}} \mathbf{B}^t \xi_{\mathbf{P}}, \tag{64}$$

where $\mathbf{e_1} = (1, 0, 0)$ and the coefficients of the matrix $\mathbf{B}$ are

$$\begin{aligned}
B_{11} &= f^3 + (1-f)^3 + f(1-f)(1-p_-)(3-p_-), \\
B_{12} &= 2f^3 p_+ + 2f^2(1-f)p_+(1-p_-), \\
B_{13} &= f^3 p_+^2, \\
B_{21} &= f(1-f)p_-(2-p_-), \\
B_{22} &= f(1-f) + (1-f)^3 + 2f(1-f)(1-p_-) + f^3(1-p_+) - f^2(1-f)p_+(1-p_-) + f^2(1-f)p_+p_-, \\
B_{23} &= f^2 p_+(1 - fp_+), \\
B_{31} &= f(1-f)p_-^2, \\
B_{32} &= 2f(1-f)(2-f)p_- + 2f^2(1-f)p_-(1-p_+), \\
B_{33} &= 1 - f + f(1 - fp_+)^2.
\end{aligned}$$

*iv. 2nd-order statistic for weights in depressed quadrant, $\langle cc^t \rangle_D$:*

The Markov process is identical, with the sole difference being the initial condition, where here $\xi_{\mathbf{D}} = \mathbf{Q_D} \xi_{\mathbf{ss}}$ with

$$\mathbf{Q_D} = \begin{pmatrix} (1-p_-)^2 & 0 & 0 \\ p_-(1-p_-) & 1-p_+ & 0 \\ p_-^2 & 2p_- & 1 \end{pmatrix}. \tag{65}$$

Then

$$\langle cc^t \rangle_D = \mathbf{e_1}^{\mathrm{T}} \mathbf{B}^t \xi_{\mathbf{D}}, \tag{66}$$

*v. Two-point statistic for synaptic weight in potentiated quadrant, $\langle c_{ij}^t c_{ij}^0 \rangle_P$:*

$$\begin{aligned}
\langle c_{ij}^t c_{ij}^0 \rangle_P &= \Pr(c_{ij}^t = 1, c_{ij}^0 = 1)_P, \\
&= \Pr(c_{ij}^t | c_{ij}^0 = 1) \Pr(c_{ij}^0 = 1)_P, \\
&= \left[\mathbf{e_1}^{\mathrm{T}} \mathbf{A^t} \mathbf{e_1}\right] \mathbf{e_1}^{\mathrm{T}} \mathbf{c_P}.
\end{aligned} \tag{67}$$

*vi. Two-point statistic for synaptic weight in depressed quadrant, $\langle c_{ij}^t c_{ij}^0 \rangle_D$:*

$$\langle c_{ij}^t c_{ij}^0 \rangle_D = \left[\mathbf{e_1}^{\mathrm{T}} \mathbf{A^t} \mathbf{e_1}\right] \mathbf{e_1}^{\mathrm{T}} \mathbf{c_D}. \tag{68}$$

#### Modeling different protocols with the Markov Process

Using this Markov process to model a simulation protocol involves using the transition matrices in the proper order. For example, to determine the state of the mean weights in the potentiated quadrant after the presentation of the tracked pattern, a number ISI of random pattern, and another repetition, is given by

$$\bar{\mathbf{c}}^{\text{final}} = \mathbf{R_p} \mathbf{A}^{\text{ISI}} \mathbf{R_p} \bar{\mathbf{c}}, \tag{69}$$

where $\bar{\mathbf{c}} = (\bar{c}, 1 - \bar{c})$. A similar operation must be performed for the second-order statistics in the potentiated quadrant, using the corresponding transition matrices, and the same for the depressed quadrant.

#### Drift in input pattern

If the input pattern itself undergoes drift from repetition to repetition, then this must be accounted for in the calculation of the correlation. We assume that if the tracked input pattern at time $t = 0$ is $\mathbf{x}$, then the $i^{\text{th}}$ input for the first repetition $x_i^1$ will be identical to that at time 0 with probability $s$ and otherwise its state will be set to active with probability $f$ or inactive with probability $1 - f$. In a slight abuse of notation we can write $\mathbf{x^1} = s\mathbf{x^0} + (1-s)\xi^1$ where $\xi$ is a random, binary vector with sparseness $f$. If this is the case, then it is straightforward to show that the $r^{\text{th}}$ input vector has correlation $s^r$ with the input vector at time $t = 0$. To model the Markov process corresponding to this choice of input vectors, for the $r^{\text{th}}$ repetition we applied the repetition transition matrix with prefactor $s^r$ and the random transition matrix with prefactor $1 - s^r$. Hence, for updating the mean weights in the potentiation vector we used $s^r\mathbf{R}_p + (1 - s^r)\mathbf{A}$.
